# VEGFA-Positive Macrophages Regulate Aqueous Humor Outflow in Aged Mice and Humans

**DOI:** 10.64898/2026.08.20.746006

**Authors:** Naoki Kiyota, Yalu Zhou, Dilip K. Deb, Guohui Ren, Tuncer Onay, Ester Reina-Torres, Hoi-Lam Li, Constance E. Runyan, Robert S. Feder, Hyunjoo J. Lee, Darryl R. Overby, Haiyan Gong, G. R. Scott Budinger, Benjamin R. Thomson, Susan E. Quaggin

**Author notes:** Correspondence to: Susan E. Quaggin, MD, 676 N. St. Clair Ave., Suite 2300, Chicago, IL 60611, USA.

## Abstract

Elevated intraocular pressure (IOP) and aging are major risk factors for primary open-angle glaucoma (POAG), but how aging affects IOP regulation remains poorly understood. IOP remains within a narrow range despite age-associated changes predicted to increase aqueous humor outflow (AHO) resistance at the interface between the trabecular meshwork and Schlemm’s canal (SC), suggesting compensatory mechanisms preserve AHO homeostasis during aging. Single-cell RNA sequencing of mouse ocular angle tissues revealed immunomodulatory transcriptional reprogramming of SC endothelial cells in older mice, while mouse and human imaging showed reduced SC size and increased peri-SC macrophage accumulation with aging. Ligand–receptor analysis predicted enhanced macrophage-to-SC VEGFA–VEGFR signaling in aged and *Tie2*-haploinsufficient mice, an independent model of vascular stress and glaucoma risk. Deletion of *Vegfa* in CX3CR1^+^ macrophages increased IOP and reduced AHO facility in 9-month-old mice, demonstrating that macrophage-derived VEGFA supports AHO homeostasis. *Tie2* haploinsufficiency recapitulated key age-associated SC niche changes, including peri-SC macrophage accumulation, whereas gene therapy boosting TIE2 activity protected wild-type mice against age-related changes. Together, these findings identify peri-SC macrophage-derived VEGFA as a compensatory mechanism maintaining AHO homeostasis during aging and vascular stress and support TIE2 activation as a therapeutic strategy to preserve SC function and IOP regulation.

## Introduction

Glaucoma is a leading cause of irreversible blindness worldwide (1). Elevated intraocular pressure (IOP), caused by increased resistance to aqueous humor outflow (AHO) (2), remains the most important modifiable risk factor for disease onset and progression, whereas aging is among the strongest nonmodifiable determinants of susceptibility (3, 4). Lowering IOP slows disease progression (5), underscoring the importance of understanding how the eye maintains IOP and resistance within a narrow physiological range. Yet how aging intersects with IOP regulation remains incompletely understood, especially because age-associated remodeling of outflow tissues does not uniformly translate into ocular hypertension. Indeed, most people do not develop elevated IOP with age (6–8). Prior experimental studies, together with imaging of healthy human eyes, suggest that age-related structural remodeling of outflow tissues occurs before overt loss of IOP regulation, raising the possibility that adaptive mechanisms preserve AHO and IOP until these responses become insufficient or are compounded by additional genetic susceptibility or environmental or mechanical stressors (9, 10).

The conventional outflow pathway consists of the trabecular meshwork (TM), Schlemm’s canal (SC), and distal outflow vessels that return aqueous humor to the venous circulation. SC is a specialized endothelial vessel with both blood vascular and lymphatic features, and its structural and functional integrity is essential for normal aqueous humor drainage and IOP homeostasis (11, 12). While considerable attention has been paid to aging-associated changes in the TM and extracellular matrix, including reduced TM cellularity, extracellular material accumulation, and tissue stiffening, the contribution of SC-specific alterations to age-related outflow dysfunction remains less well defined (9, 13–15).

In other vascular beds, aging is associated with endothelial activation and enhanced immune-endothelial interactions, often involving NF-κB-centered programs and increased leukocyte recruitment (16, 17). At the same time, specialized macrophage subsets can support tissue homeostasis by maintaining vascular integrity and restraining age-related inflammation (18, 19). Consistent with this broader role for immune cells in vascular function, recent work demonstrated that resident tissue macrophages in the TM/SC support IOP and outflow homeostasis (20). However, while these observations suggest inflammatory cells may modulate IOP, whether these cells play a role in the aged SC and eye, how such cells might relate to glaucoma susceptibility, and what molecular pathways are responsible for macrophage-mediated regulation of IOP remain unknown.

Here we tested the hypothesis that compensatory age-related changes in outflow tissues of the eye maintain IOP homeostasis despite deleterious structural and mechanical changes that would otherwise lead to increased IOP, and that genetic susceptibility to glaucoma may engage overlapping adaptive responses. Using a combination of single-cell RNA sequencing, imaging in human and mouse eyes, and functional analyses in mouse models, we found that *VEGFA*-expressing CX3CR1^+^ macrophages accumulate around the SC in older mice and humans and in young *Tie2*-haploinsufficient mice, a genetic model of glaucoma risk, and that these macrophages modulate IOP. TM-targeted adeno-associated virus (AAV) gene delivery of the TIE2 ligand, ANGPT1, improves outflow facility and lowers IOP in *Tie2*-deficient and wild-type eyes, and reduces age-related adaptive responses. These findings provide a new model for age-related adaptations that modulate AHO and identify an SC-targeted approach to slow age-related changes in the SC–TM niche.

## Results

### Aging induces inflammatory transcriptional reprogramming of Schlemm’s canal and local immune cell accumulation

To define how aging reshapes the conventional outflow pathway at the molecular level, we performed single-cell RNA sequencing (scRNA-seq) on iridocorneal angle tissues from the eyes of young (6-week-old) and older (15-month-old) mice (Fig. 1A). These timepoints were selected based on human epidemiologic data showing that POAG prevalence and incidence rise substantially from the age of 50 and onwards (21), with 44% of cases occurring between 55 and 74 years of age (22). scRNA-seq tissue processing followed our previously published workflow (23). Each sample consisted of six pooled eyes (3 mice), generating two independent samples per age group (n = 2 samples per group). We performed unsupervised clustering and annotated cell identities in accordance with our prior single-cell atlas (24). Marker expression patterns supporting these annotations are shown in Supplementary Fig. S1. Uniform manifold approximation and projection (UMAP) resolved major anterior segment cell populations, including trabecular meshwork (TM; *Myoc*^+^, *Chil1*^+^), multiple immune cell populations, and stromal cell populations, and identified a discrete SC endothelial cluster (*Cdh5*^+^, *Prox1*^+^, *Lyve1*^−^) (Fig. 1B, arrow). To identify age-dependent transcriptional changes, we extracted the SC cluster and performed differential expression analysis comparing young and older SC cells. Enrichr pathway enrichment analysis using the top 200 upregulated genes showed enrichment of inflammatory programs in older SC cells, including “inflammatory response” and “TNF alpha signaling via NF-κB” (Fig. 1C). This interpretation is consistent with evidence that vascular aging is accompanied by a pro-inflammatory endothelial phenotype in which multiple inflammatory pathways converge on NF-κB (16, 17). Representative genes contributing to the “inflammatory response” and “TNF alpha signaling via NF-κB” gene sets are highlighted in Fig. 1D. These analyses indicated that older SC cells exhibited increased expression of inflammatory and immune-interaction genes, including *Icam1* and *Cx3cl1*, alongside other genes within these inflammatory/TNFα-associated signatures. We therefore examined whether this transcriptional state was accompanied by tissue-level remodeling and local immune accumulation. Eyes were collected from wild-type mice across an age series (6 weeks, 3 months, 6 months, 9 months, 12 months, and 15 months), and anterior segment flat mounts were prepared to evaluate SC morphology and immune cell populations. Immunostaining revealed an age-dependent increase in CD45^+^ cells in the SC region, hereafter referred to as the peri-SC region (Fig. 1E). SC area (×10^5^ μm^2^ per 20× image) progressively decreased with age (one-way ANOVA; P = 0.0001), with significant reductions at 12 months (1.150 ± 0.051; P = 0.0336) and 15 months (1.118 ± 0.040; P = 0.0065) compared with 6-week controls (1.299 ± 0.026) by Dunnett’s multiple-comparisons test (Fig. 1F). In parallel, CD45^+^ area per SC area (%) increased with age (one-way ANOVA; P = 0.0053), with significant elevations versus 6-week controls at 9 months (16.73 ± 0.65; P = 0.0236), 12 months (16.87 ± 0.49; P = 0.0142), and 15 months (17.47 ± 0.65; P = 0.0012) by Dunnett’s multiple-comparisons test (Fig. 1G).

**Figure 1.**
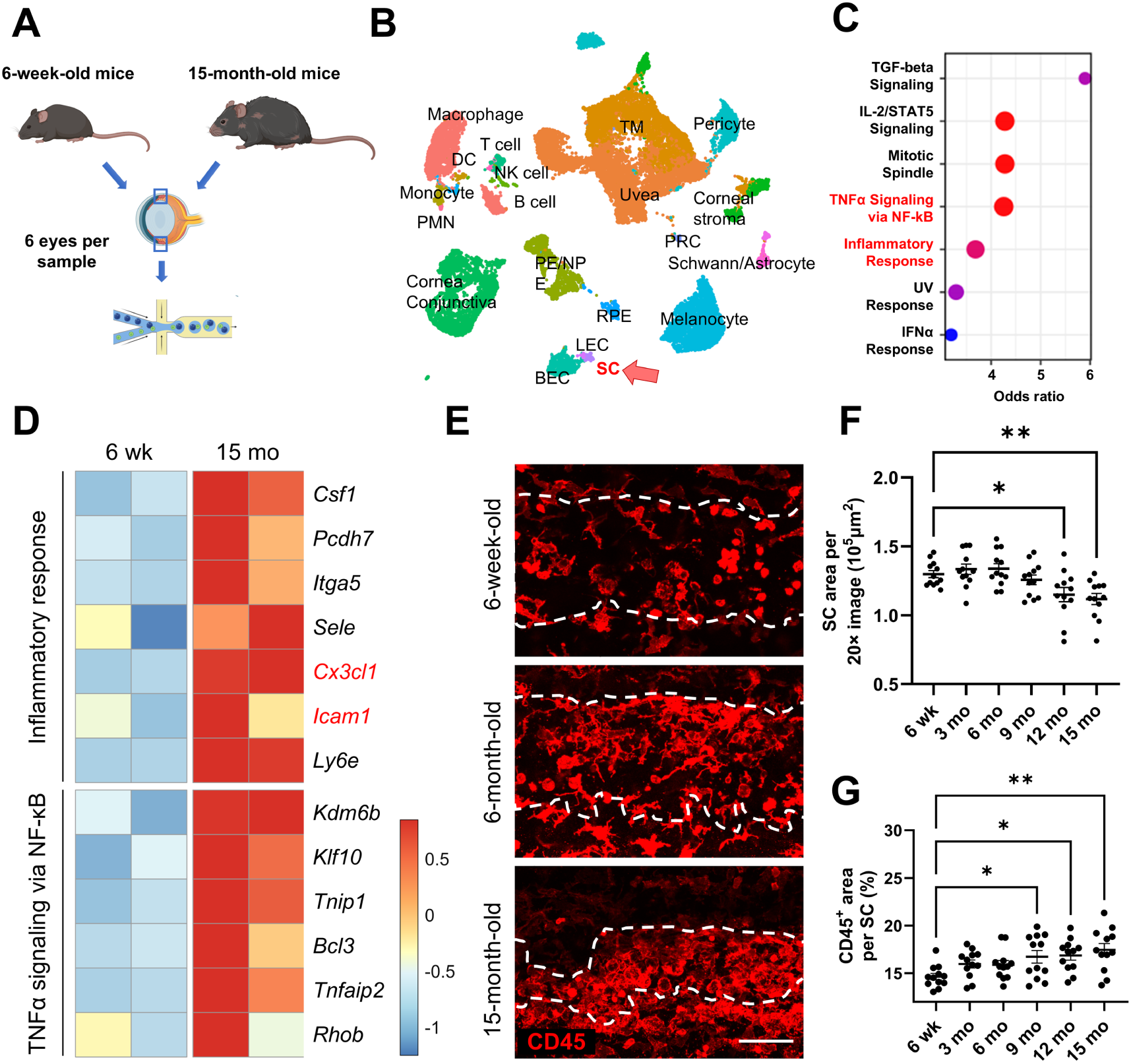
Aging induces inflammatory transcriptional reprogramming of Schlemm’s canal and promotes local immune accumulation. (A) Experimental schematic for single-cell RNA sequencing (scRNA-seq) of limbal/iridocorneal angle tissues from young (6-week-old) and older (15-month-old) mice. Tissues were pooled as six eyes per sample, generating two independent samples per age group. (B) Uniform manifold approximation and projection (UMAP) of anterior segment cell populations identified by scRNA-seq. The arrow indicates the Schlemm’s canal (SC) endothelial cluster. (C) Enrichr pathway enrichment analysis using the top 200 genes upregulated in older SC cells, highlighting inflammatory programs including inflammatory response and TNF alpha signaling via NF-κB. (D) Heatmap showing representative genes contributing to the indicated inflammatory gene sets in SC cells from young (6 wk) and older (15 mo) mice; each column represents one biological replicate. (E) Representative CD45 immunostaining images of anterior segment flat mounts from 6-week-old, 6-month-old, and 15-month-old mice, showing an age-related increase in CD45^+^ signal within and adjacent to the SC region. Dashed lines outline the SC region. Scale bar, 100 μm. (F) Quantification of SC area across the indicated ages. SC area progressively decreased with age, with significant reductions at 12 and 15 months compared with 6-week-old controls. (G) Quantification of CD45^+^ area normalized to SC area across the indicated ages. Normalized CD45^+^ area per SC area increased with age, with significant elevations at 9, 12, and 15 months compared with 6-week-old controls. In (F) and (G), each dot represents one mouse, with values averaged between both eyes (n = 12 mice per age group). Data are presented as mean ± SEM. One-way ANOVA followed by Dunnett’s multiple-comparisons test against 6-week-old controls was used in (F) and (G). Only significant comparisons are shown. *P < 0.05, **P < 0.01.

### Aging is associated with peri-SC macrophage accumulation and reduced SC dimensions in mice and humans

Immunostaining revealed that the majority of CD45^+^ leukocytes we identified in the SC region co-expressed IBA1, a well-validated macrophage marker (IBA1^+^/CD45^+^, 6-week: 85.62 ± 4.23% vs 15-month: 89.76 ± 2.56%, mean ± SEM; P = 0.481, Student’s two-tailed unpaired t-test, Supplementary Fig. S2). Direct comparison of macrophage accumulation at 6 weeks and 15 months revealed an increased IBA1^+^ macrophage density within and adjacent to the SC in older mice (IBA1^+^ cells per 10^4^ μm^2^ SC: 6-week: 8.914 ± 0.680 vs 15-month: 13.07 ± 0.46; P = 0.0012, two-tailed unpaired t-test, Fig. 2A,C). Consistent with the age-series analysis, SC area was reduced in older mice (6-week: 1.299 ± 0.026 vs 15-month: 1.118 ± 0.040 ×10^5^ μm^2^ per 20× image; P = 0.0010, two-tailed unpaired t-test, Fig. 2A,B). To determine if similar age-associated changes occurred in human eyes, we analyzed immunostained cryosections of corneal rim tissues containing SC and trabecular meshwork (donor age and death-to-preservation interval are summarized in Supplementary Table S1). SC circumferential length was shorter in donors aged ≥ 50 years than in those aged < 50 years (526.0 ± 71.6 vs 857.5 ± 88.0 μm; P = 0.0281, Student’s two-tailed unpaired t-test), whereas IBA1^+^ macrophage density normalized to circumferential SC length was higher (3.129 ± 0.382 vs 1.606 ± 0.146 cells per 100 μm SC; P = 0.0263, Student’s two-tailed unpaired t-test) (Fig. 2D–F). Thus, across both mice and humans, aging was associated with greater peri-SC macrophage accumulation and reduced SC dimensions, reflected by decreased SC area in mice and shorter circumferential length in humans.

**Figure 2.**
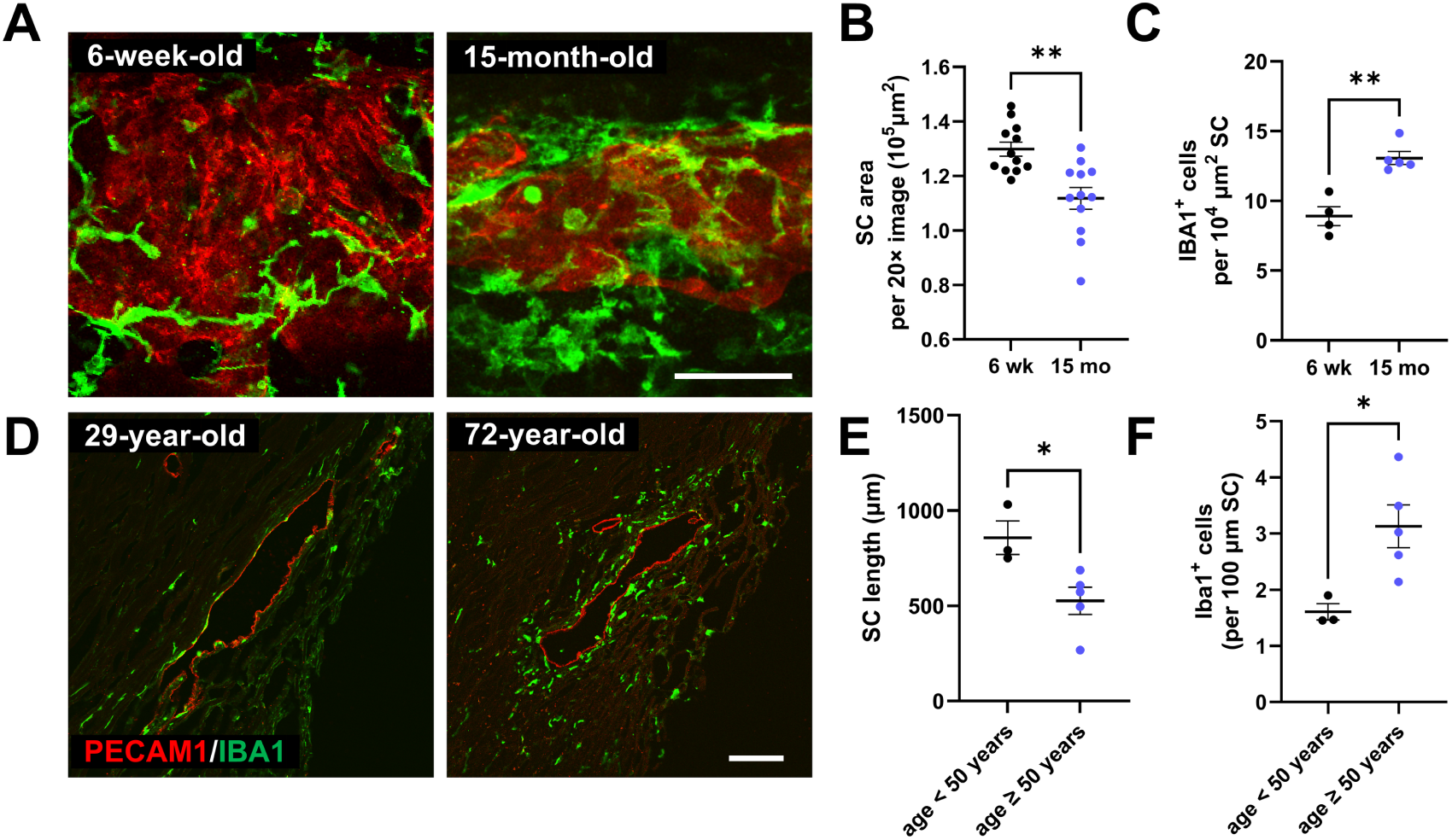
Aging is associated with peri-SC macrophage accumulation and reduced Schlemm’s canal dimensions in mice and humans. (A) Representative PECAM1/IBA1 immunostaining images of anterior segment flat mounts from 6-week-old and 15-month-old mice. Older mice show reduced Schlemm’s canal (SC) area and increased accumulation of peri-SC IBA1^+^ macrophages. Scale bar, 100 μm. (B) Quantification of SC area in young and older mouse eyes. (C) Quantification of IBA1^+^ macrophage density normalized to SC area in young and older mouse eyes. (D) Representative PECAM1/IBA1 immunostaining images of human limbal sections from a 29-year-old donor and a 72-year-old donor, showing shorter circumferential SC length and greater peri-SC macrophage accumulation in the older donor. Scale bar, 100 μm. (E) Quantification of circumferential SC length in human donor samples stratified by age (<50 years and ≥50 years). (F) Quantification of IBA1^+^ macrophages normalized to circumferential SC length in human donor samples stratified by age. Data are presented as mean ± SEM. Each dot represents an individual mouse in (B) and (C), with values averaged between both eyes, or an individual human donor (one eye per donor) in (E) and (F). Two-tailed unpaired t-tests were used for comparisons in (B) and (C). Student’s two-tailed unpaired t-tests were used in (E) and (F). *P < 0.05, **P < 0.01.

### Aging enhances SC–macrophage crosstalk via the CX3CL1–CX3CR1 axis and promotes peri-SC accumulation of CX3CR1^+^ macrophages

Given the age-associated increase in peri-SC macrophages, we revisited our single-cell analysis to identify macrophage populations that might preferentially communicate with SC. Sub-clustering of myeloid cells resolved monocytes (Mono; *Ace*, *Spn*, and *Ly6c2*) and five *Aif1*-expressing macrophage populations (Mac1–Mac5, Fig. 3A). Of these, Mac1 exhibited a *Cx3cr1*-high, MHC class II–high profile, with low expression of common markers of tissue resident macrophages including *Mrc1*, *Pf4*, and *Gas6*. Mac2 exhibited an inflammatory profile, expressing *Ccl5* and high levels of MHC class II–associated genes. Mac4 and Mac5 expressed markers associated with tissue-resident macrophages, including *Mrc1* (encoding CD206), *Lyve1*, and *Pf4* (25, 26). In addition to these markers, Mac5 expressed *Cx3cr1* and lower levels of *Lyve1* than Mac4, with *Lyve1* detected in 55.6% of Mac4 cells (70/126), compared with 30.1% of Mac5 cells (126/419, Fig. 3B).

**Figure 3.**
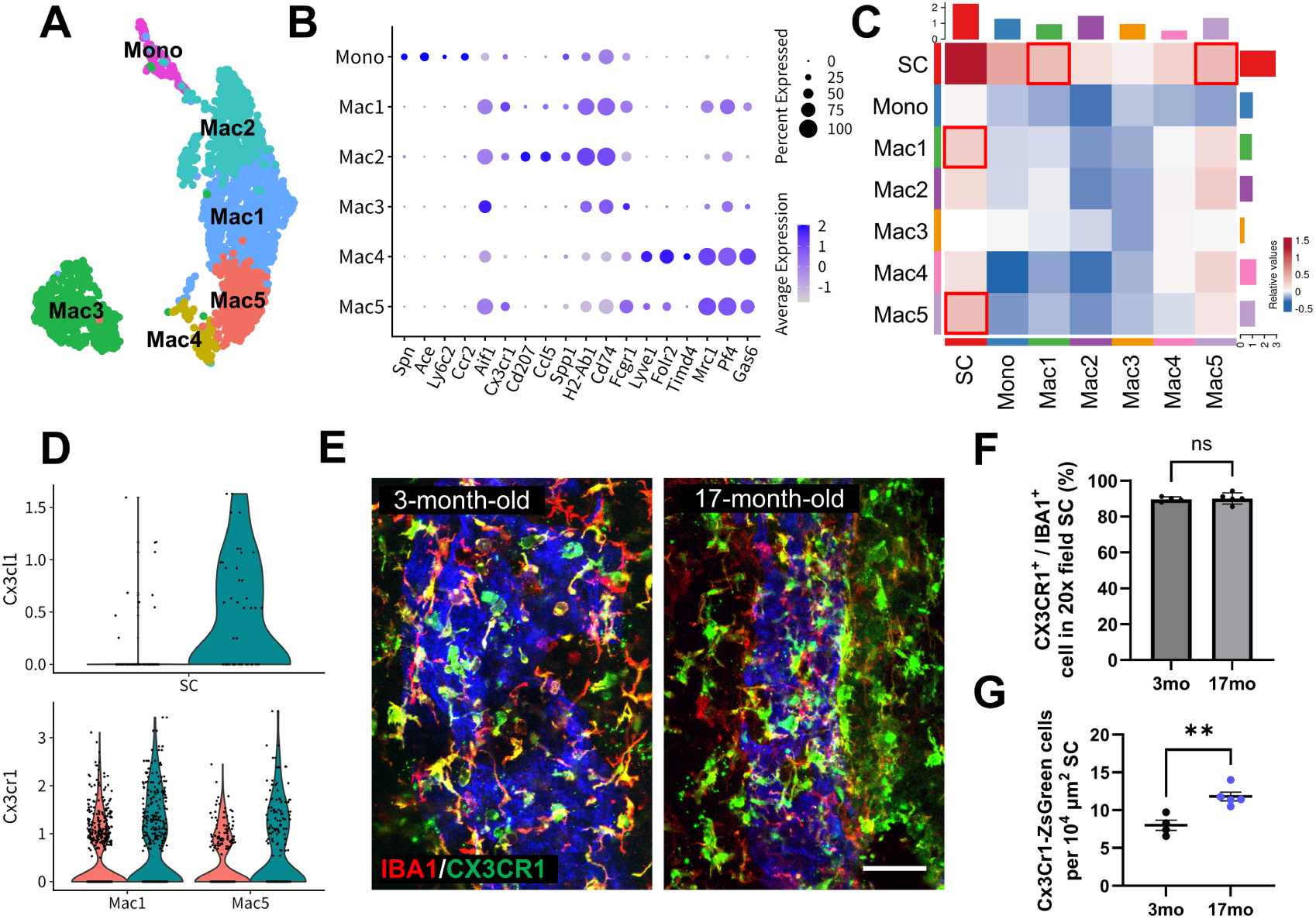
Aging enhances SC–macrophage crosstalk via the CX3CL1–CX3CR1 axis and promotes peri-SC accumulation of CX3CR1^+^ macrophages. (A) UMAP of the myeloid compartment showing monocytes (Mono) and five macrophage subclusters (Mac1–Mac5). (B) Dot plot showing marker expression across monocytes and macrophage subclusters, including *Cx3cr1* and *Lyve1* expression in Mac1 and Mac5. (C) CellChat interaction-w*eight* heatmap showing bidirectional communicat*ion be*tween Schlemm’s canal (SC) and myeloid subclusters. Red boxes highlight the strongest SC–macrophage interactions, observed with Mac1 and Mac5. (D) Violin plots showing Cx3cl1 expression in SC endothelial cells and Cx3cr1 expression in Mac1 and Mac5 from young and older eyes. (E) Representative IBA1/CX3CR1 immunostaining images of the SC region from 3-month-old and 17-month-old *Cx3cr1*-CreERT2; ZsGreen reporter mice, showing increased accumulation of CX3CR1^+^ macrophages with aging. Scale bar, 100 μm. (F) Quantification of the fraction of CX3CR1^+^ cells among IBA1^+^ macrophages in the SC region. (G) Quantification of CX3CR1^+^ (ZsGreen^+^) macrophage density in the SC region. Data are presented as mean ± SEM. In (F) and (G), each dot represents one eye from an individual mouse. Student’s two-tailed unpaired t-test was used in (F), and a two-tailed unpaired t-test was used in (G). ns, not significant; **P < 0.01.

Following subclustering, CellChat analysis was used to predict bidirectional communication between macrophage populations and the SC endothelium, revealing that this crosstalk was most prominent for Mac1 and Mac5 compared with the other macrophage subclusters (Fig. 3C), and that the CX3CL1–CX3CR1 pathway was a major axis of crosstalk between these cell types (Supplementary Fig. S3C). Older SC endothelial cells showed higher expression of *Cx3cl1*, which encodes the ligand and adhesion molecule for CX3CR1, whereas *Cx3cr1* was expressed by Mac1 and Mac5, supporting potential CX3CL1–CX3CR1 interactions between SC and these two macrophage populations (Fig. 3D). Based on these findings, we hypothesized that macrophages accumulating in the peri-SC niche with age and predicted to communicate with SC are predominantly CX3CR1^+^LYVE1-low/negative. To test this prediction in vivo and identify Cx3cr1^+^ macrophages, we used *Cx3cr1*-CreERT2; ZsGreen reporter mice. Tamoxifen was administered by oral gavage (10 mg/100 µL) twice, 7 and 2 days before eye collection, and eyes were collected at 3 or 17 months of age.

Immunostaining showed that nearly all peri-SC IBA1^+^ macrophages were CX3CR1^+^ at both ages, with no significant age-dependent change in this proportion (Fig. 3E,F). CX3CR1^+^LYVE1^+^ macrophages were rare in the SC region and represented 2.87% ± 0.42% of peri-SC CX3CR1^+^ macrophages (mean ± SEM, n = 5). Among the few LYVE1^+^ cells observed in the SC region, most were CX3CR1^+^, whereas LYVE1^+^ cells in distal outflow pathway blood endothelial cell/lymphatic endothelial cell (BEC/LEC) regions showed more heterogeneous CX3CR1 expression (Supplementary Fig. S3A,B). The density of CX3CR1^+^ peri-SC macrophages increased significantly with age (3 months: 8.01 ± 0.67 vs. 17 months: 11.81 ± 0.59 CX3CR1-ZsGreen^+^ cells per 10⁴ μm² SC, mean ± SEM; *P* = 0.0037, two-tailed unpaired t test; Fig. 3E,G). These findings indicate that the peri-SC macrophage population that increases with age exhibits a predominantly CX3CR1^+^ and LYVE1-low/negative phenotype. This phenotype is consistent with Mac1 and may also correspond to the LYVE1-low/negative subset within Mac5.

### VEGFA is expressed by peri-SC macrophages in older mice and humans

To identify molecular pathways that might underlie adaptive or maladaptive responses in the aging SC niche, we analyzed scRNA-seq data and performed CellChat analysis to identify SC-macrophage crosstalk pathways that were altered by aging. This analysis predicted significantly increased Mac1- and Mac5-to-SC *Vegfa*–Vegfr signaling in older compared with young eyes, including *Vegfa*–*Vegfr2*, *Vegfa*–*Vegfr1r2*, and *Vegfa*–*Vegfr1* interactions (Fig. 4A). To visualize the expression patterns underlying this interaction-level result, we plotted *Vegfa* expression in Mac1 and Mac5 and *Kdr* expression in SC endothelial cells; these distributions were consistent with the CellChat prediction (Fig. 4B). As VEGFA is known to regulate outflow facility (27), we speculated that macrophage-to-SC VEGFA signaling may represent one pathway through which peri-SC macrophages support outflow function. To validate the age-associated increase in macrophage *Vegfa* expression in vivo, we performed *Vegfa* RNAscope in combination with CD68 immunolabeling around the SC in mouse eyes. In mice, *Vegfa*-expressing CD68^+^ peri-SC macrophages were more frequently detected in older (15-month-old) than in young (6-week-old) eyes (Fig. 4C), and quantification confirmed a significant increase in *Vegfa*-expressing CD68^+^ cells per 100 μm SC in older mice (Fig. 4D). In human eyes, *VEGFA*-expressing IBA1^+^ macrophages were also detected around the SC (Fig. 4E), and the fraction of *VEGFA*-expressing cells among peri-SC IBA1^+^ macrophages was positively correlated with donor age (Pearson *r* = 0.84, *P* = 0.009; Fig. 4F; donor information summarized in Supplementary Table S2).

**Figure 4.**
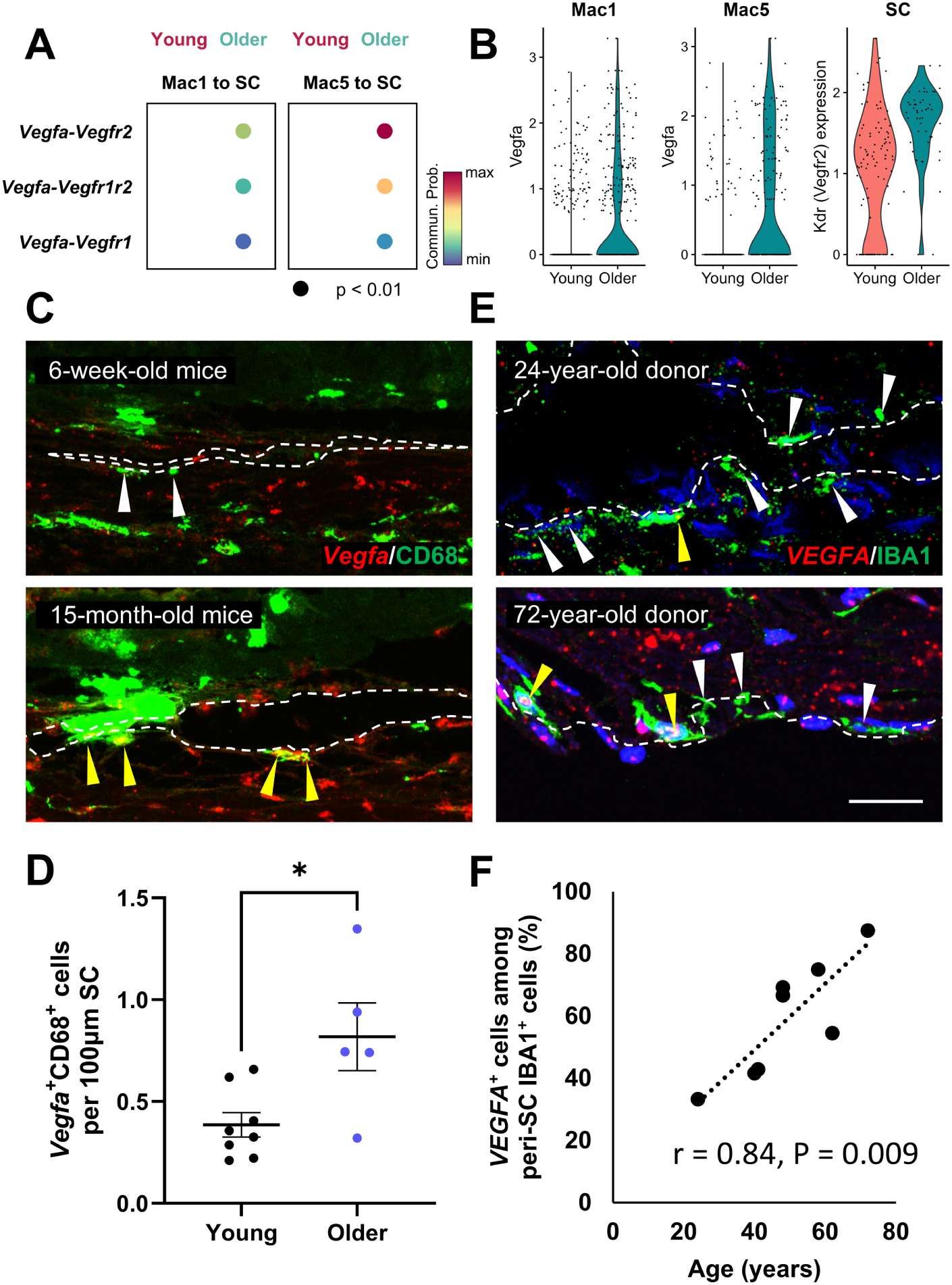
Aging increases macrophage-associated *Vegfa*/VEGFA expression around the SC. (A) Dot plot showing selected Mac1- and Mac5-to-SC *Vegfa–Vegfr* interactions predicted by CellChat analysis in the aging dataset. Dot color indicates communication *p*robability; all displayed interact*ions* had P < 0.01. (B) Violin plot*s sho*wing Vegfa expression in Mac1 and Mac5 and Kdr expression in SC endothelial cells from young and older eyes. (C) Representative Vegfa RNAscope/CD68 immunolabeling *image*s of the SC region in 6-week-old and 15-mont*h-old* mouse eyes. Dashed lines outline the SC region. White arrowheads *indi*cate CD68^+^ macrophages without detectable *Vegfa* signal, whereas yellow arrowheads indicate *Vegfa*-expressing CD68^+^ macrophages around the SC. (D) Quantification of *Vegfa*-expressing CD68^+^ cells per 100 μm SC in young and older mouse eyes. (E) Representative VEGFA/IBA1 images of human limbal sections from a 24-year-old donor and a 72-year-old donor. Dashed lines outline the SC region. White arrowheads indicate IBA1^+^ macrophages without detectable VEGFA signal, whereas yellow arrowheads indicate *VEGFA*-expressing IBA1^+^ macrophages around the SC. (F) Correlation between donor age and the fraction of *VEGFA*-expressing macrophages among total peri-SC IBA1^+^ macrophages. Scale bar, 50 μm. Data are presented as mean ± SEM. In (D), each dot represents one mouse, with values averaged between both eyes. In (F), each dot represents one human donor (one eye per donor). A two-tailed unpaired Student’s t-test was used in (D). Pearson correlation analysis was used in (F). *P < 0.05.

### Peri-SC macrophages are identified in regions of enhanced AH outflow in aged mice

Previous tracer-based perfusion studies and aqueous angiography have established that conventional outflow is segmental, with circumferentially distinct high-flow and low-flow regions along SC (28, 29). To examine how aging-associated peri-SC macrophage accumulation relates to these segmental flow domains, we labeled outflow with a fluorescent tracer and immunostained anterior segment flat mounts for IBA1 (Fig. 5A). In young eyes, IBA1 signals were sparse (stars) and tracer and IBA1 signals were negatively correlated within the SC region (Young: −0.211 ± 0.022, n = 6 mice; Fig. 5A,B), consistent with the broader observation that low-flow/low-shear vascular domains preferentially support leukocyte recruitment (30). By contrast, in older eyes, IBA1^+^ cells were more frequently observed near tracer-positive regions including high-flow domains (white arrow; Fig. 5A), supporting a model whereby accumulating macrophages may contribute to outflow homeostasis. The identification that the peri-SC macrophages in older eyes express Vegfa, which is known to enhance outflow, supports this model. Finally, the negative correlation between tracer and IBA1 signals was significantly weaker in older eyes (Older: −0.137 ± 0.018, n = 6 mice; P = 0.0276; Fig. 5B). IOP did not show a clear age-dependent change across this age range (Supplementary Fig. S4), consistent with maintained pressure homeostasis despite age-associated SC remodeling.

**Figure 5.**
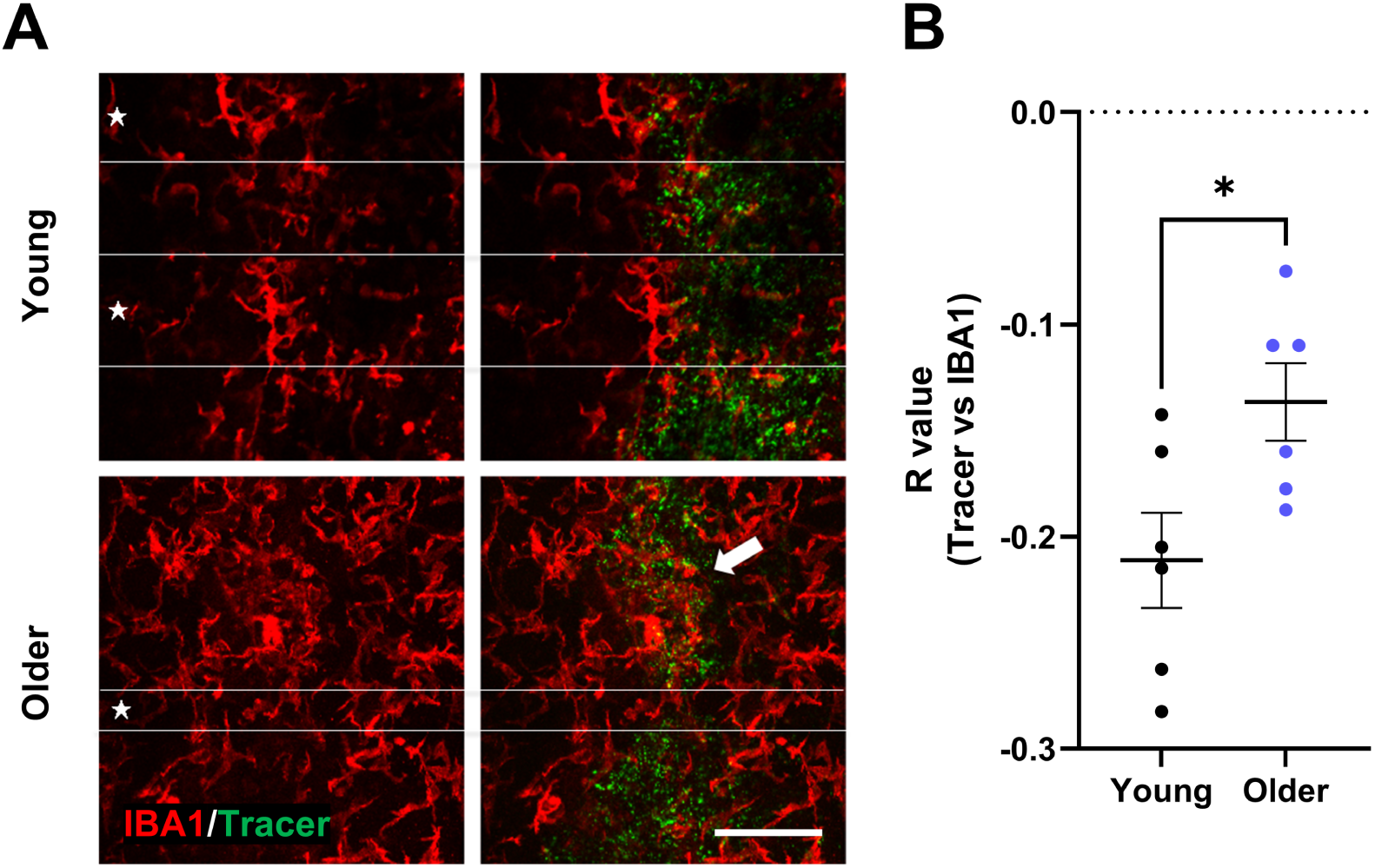
Aging reduces the preferential localization of peri-SC macrophages to low-flow regions. (A) Representative anterior segment flat-mount images labeled with fluorescent tracer and immunostained for IBA1 in young and older eyes. In young eyes, tracer-enriched regions show relatively sparse IBA1 signal (stars), whereas in older eyes, IBA1^+^ cells are more frequently observed adjacent to tracer-positive regions, and occasional overlap between macrophage accumulation and tracer-enriched domains is apparent (white arrow). (B) Quantification of the Pearson correlation coefficient between tracer and IBA1 signals within the Schlemm’s canal (SC) region. Scale bar, 100 μm. Data are presented as mean ± SEM. Each dot represents one mouse, with values averaged across eight quadrants from both eyes. Statistical analysis was performed using a two-tailed unpaired Student’s t-test. *P < 0.05.

### *Vegfa* deletion in CX3CR1^+^ macrophages increases IOP and reduces outflow facility in aged mice

To test whether macrophage-derived VEGFA in the eye plays a functional role in AHO during aging, we conditionally deleted *Vegfa* from *Cx3cr1*^+^ macrophages by crossing mice carrying a *Vegfa* floxed allele (Supplementary Fig. S5) with *Cx3cr1*-CreERT2 mice to obtain *Cx3cr1*-CreERT2; *Vegfa*^fl/fl^ mice (hereafter, “CKO”). Tamoxifen was administered intraperitoneally at 6 weeks of age (75 mg/kg/day for 3 days; Fig. 6A), and the change in IOP relative to pretreatment baseline (ΔIOP) was assessed over time. ΔIOP did not differ between *Vegfa* WT and *Vegfa* CKO eyes at 3 or 4 months of age but was significantly increased in *Vegfa* CKO eyes at 6 and 9 months of age (Fig. 6B,C). At 9 months, *Vegfa* CKO eyes also exhibited reduced outflow facility compared with *Vegfa* WT eyes (Fig. 6D). To confirm deletion of *Vegfa* in peri-SC macrophages in the 9-month cohort, we performed *Vegfa* RNAscope combined with IBA1 immunostaining and observed a reduced fraction of *Vegfa*-expressing IBA1^+^ macrophages around the SC in *Vegfa* CKO eyes (Supplementary Fig. S6). Because *Vegfa* deletion was induced with a pulse of tamoxifen at 6 weeks of age, whereas IOP elevation first became evident at 6 months and persisted through 9 months, these findings suggest that VEGFA produced by long-lived CX3CR1-lineage cells contributes to maintenance of outflow homeostasis with age. Circulating monocytes express CX3CR1 but have a short lifespan, such that tamoxifen-induced recombination in this population is transient as labeled monocytes are replaced by newly generated, unrecombined cells (31, 32). In contrast, recombination can persist in long-lived tissue-resident CX3CR1^+^ macrophages (31). Thus, the emergence of the phenotype months after the tamoxifen pulse is most consistent with a functional requirement for VEGFA derived from long-lived tissue-resident macrophages.

**Figure 6.**
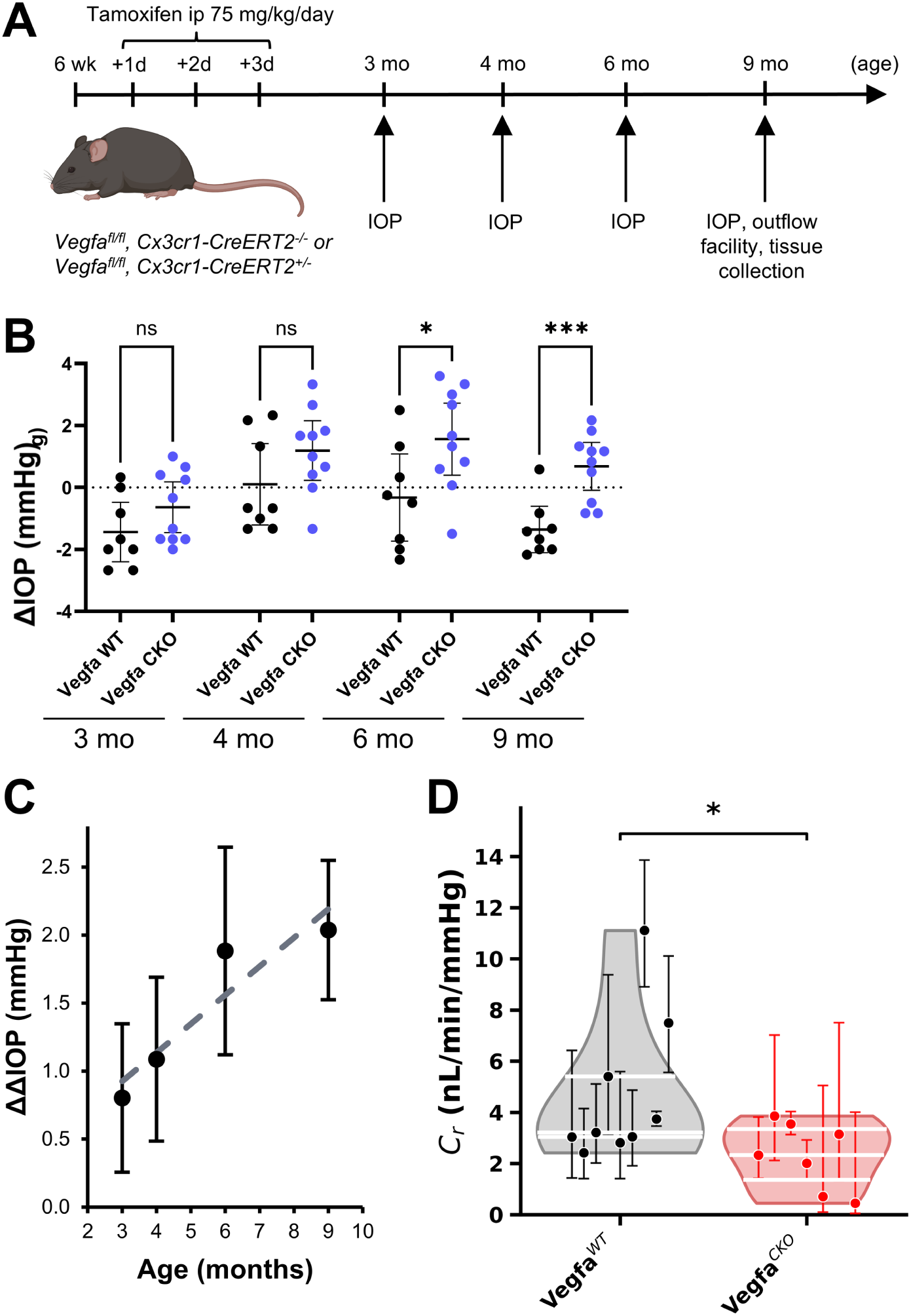
*Vegfa* deletion in CX3CR1^+^ macrophages increases IOP and reduces outflow facility. (A) Experimental timeline. *Vegfa*^fl/fl^; *Cx3cr1*-CreERT2^+/−^ mice (*Vegfa* CKO) or *Vegfa*^fl/fl^; *Cx3cr1*-CreERT2^−/−^ littermate controls (*Vegfa* WT) received intraperitoneal tamoxifen (75 mg/kg/day) for 3 consecutive days beginning at 6 weeks of age. IOP was measured at 3, 4, 6, and 9 months of age, and outflow facility and anterior segments were analyzed at 9 months. (B) Change in IOP relative to pretreatment baseline (ΔIOP) in *Vegfa* WT and *Vegfa* CKO eyes at each time point. ΔIOP did not differ between groups at 3 or 4 months of age, but was significantly increased in *Vegfa* CKO eyes at 6 and 9 months. (C) Age-dependent difference in ΔIOP between *Vegfa* CKO and *Vegfa* WT eyes, plotted as the CKO-minus-WT mean difference at each time point. (D) Quantification of conventional outflow facility (Cr) in 9-month-old *Vegfa* WT and *Vegfa* CKO eyes. In (B), each dot represents one mouse, with ΔIOP calculated from IOP values averaged between both eyes, and bars indicate mean with 95% confidence interval. In (C), points indicate CKO-minus-WT mean difference with SEM. In (D), each dot represents one eye from an individual mouse; raw Cr values are shown with asymmetric confidence intervals from individual iPerfusion fits and violin distributions. Statistical analysis in (B) used row-wise Student’s two-tailed unpaired t-tests. Statistical analysis in (D) was performed using Student’s two-tailed unpaired t-test on ln(Cr). ns, not significant; *P < 0.05; ***P < 0.001.

### Tie2 haploinsufficiency is a genetic model for glaucoma risk and converges on age-associated SC niche remodeling

To further test the hypothesis that overt ocular hypertension develops when age-related adaptive mechanisms are exceeded by additional genetic, mechanical, or environmental stressors, we studied mice carrying a *Tie2*-deficient allele. TIE2 signaling is required for SC development and adult maintenance, and its impairment is associated with ocular hypertension and glaucoma in mice and humans (33, 34). *Tie2* heterozygous-null mice (*Tie2*^+/−^) were generated through EIIa-Cre-mediated germline recombination and compared with *Tie2*^+/+^ littermate controls (Fig. 7A). At 6 weeks of age, IOP was modestly higher in *Tie2*^+/−^ mice than in *Tie2*^+/+^ controls (*Tie2*^+/+^: 10.23 ± 0.21 mmHg vs. *Tie2*^+/−^: 11.66 ± 0.34 mmHg; adjusted *P* = 0.0089), consistent with the previously reported ocular hypertensive phenotype of *Tie2* haploinsufficiency (33). By 9 months, IOP remained relatively stable in *Tie2*^+/+^ mice (10.59 ± 0.40 mmHg) but was further elevated in *Tie2*^+/−^ mice (13.92 ± 0.42 mmHg; adjusted *P* < 0.0001) (Fig. 7B). Two-way ANOVA revealed a significant genotype × age interaction (*P* = 0.0076), indicating that the age-dependent change in IOP differed by genotype and was greater in *Tie2*-haploinsufficient mice (main effects: genotype, *P* < 0.0001; age, *P* = 0.0003). Šídák’s multiple-comparisons test confirmed significant genotype differences at both ages, supporting the ‘two or multiple hit’ hypothesis that genetic susceptibility amplifies age-dependent glaucoma risk.

**Figure 7.**
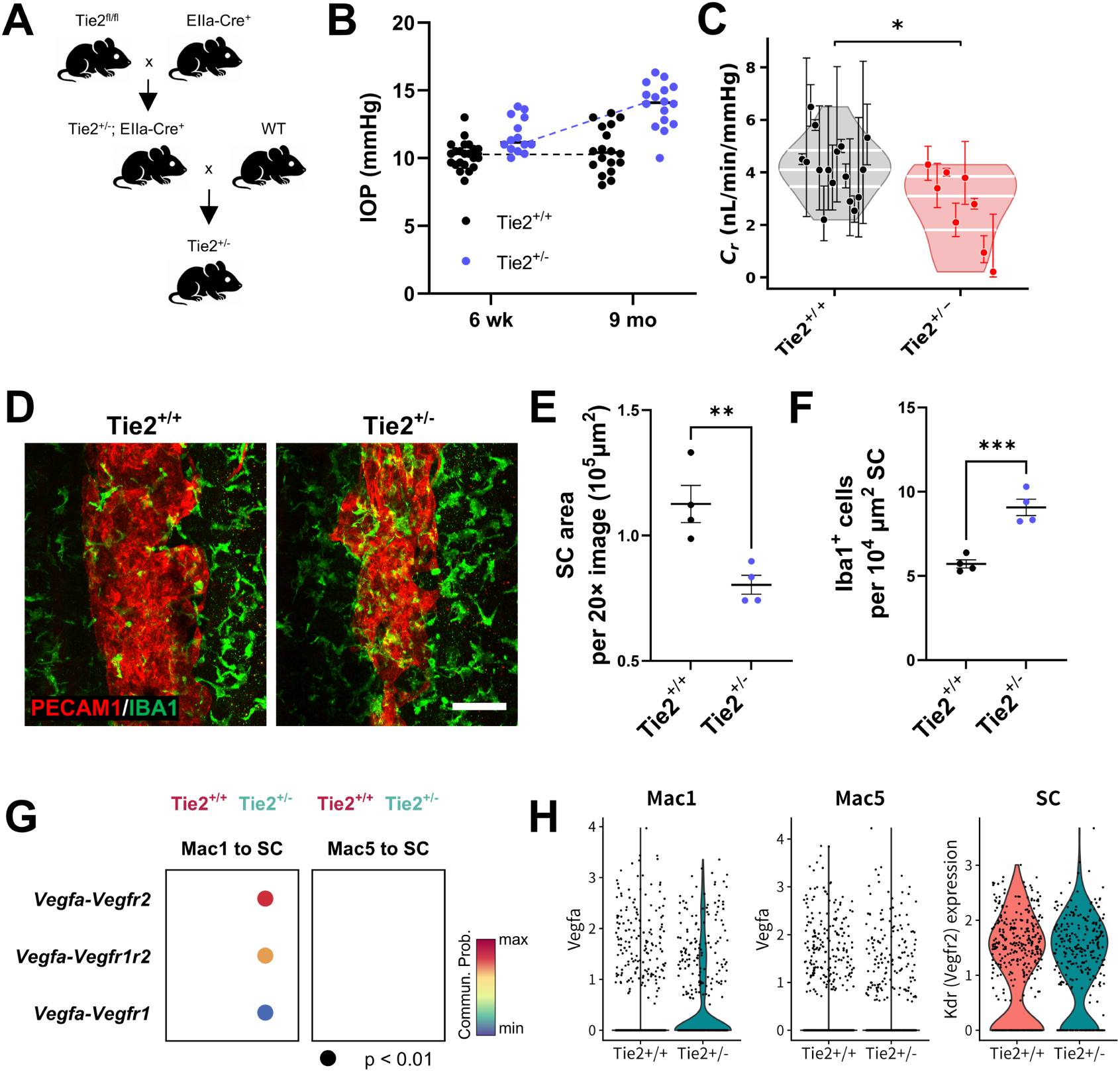
*Tie2* haploinsufficiency reproduces reduced SC area and peri-SC macrophage accumulation seen with aging. (A) Breeding scheme used to generate *Tie2* heterozygous-null mice (*Tie2*^+/−^) via EIIa-Cre-mediated germline recombination. (B) Intraocular pressure (IOP) measurements in *Tie2*^+/+^ and *Tie2*^+/−^ mice at 6 weeks (6 wk) and 9 months (9 mo) of age. (C) Quantifica*tion* of outf*low* facility (Cr) in 9-month-old Tie2^+/+^ and Tie2^+/−^ eyes. *(D)* Representative PECAM1/IBA1 immunostaining images of anterior segment flat mounts from 6-week-old Tie2^+/+^ and Tie2^+/−^ mice. Scale bar, 100 μm. (E) Qua*ntifi*c*ation* of Schlemm’s canal (SC) area. (F) Quantification of peri-SC IB*A1^+^* macrophage density normalized t*o SC* a*rea.* (G) Dot plots showing selected *Mac1*- and Mac5-to-SC *Vegfa–Vegfr* interactions predicted by CellChat analysis in the 6-week-old Tie2 scRNA-seq dataset. Mac1-to-SC Vegfa–Vegfr interactions w*ere d*etected in Tie2-haploinsufficient *bu*t not wild-type eyes, whereas corresponding Mac5-to-S*C in*teractions were not detected in either genotype. (H) Violin plots showing Vegfa expression in Mac1 and Mac5 and Kdr expression in SC endothelial cells in wild-type and Tie2-haploinsufficient eyes. Data are *p*resented as mean ± SEM. Each dot represents one mouse, with IOP averaged between both eyes, in (B), or one eye from an individual mouse in (C), (E), and (F). In (B), two-way ANOVA followed by Šídák’s multiple-comparisons test was used; adjusted P = 0.0089 at 6 weeks and adjusted P < 0.0001 at 9 months. In (C), statistical analysis was performed using a two-tailed unpaired Student’s t-test on ln(C_r_). In (E) and (F), two-tailed unpaired Student’s t-tests were used. Dot color in (G) indicates communication probability; all displayed interactions had P < 0.01. *P < 0.05, **P < 0.01, ***P < 0.001.

At 9 months, outflow facility (Cr) was also significantly reduced in *Tie2*^+/−^ eyes compared with *Tie2*^+/+^ controls (*Tie2*^+/+^: 4.17 ± 0.29 vs. *Tie2*^+/−^: 2.70 ± 0.53 nL/min/mmHg; n = 16 and 8 eyes, respectively; unpaired two-tailed Student’s *t* test on ln(Cr), *P* = 0.0186) (Fig. 7C). While we predicted increased glaucoma risk and elevated IOP in older mice carrying a second insult such as a genetic susceptibility, somewhat unexpectedly, we found that Tie2 haploinsufficiency produced SC niche remodeling similar to that observed with aging, characterized by accumulation of Cx3cr1^+^ macrophages.

At 6 weeks of age, PECAM1/IBA1 immunostaining of anterior segment flat mounts showed reduced SC area and increased peri-SC macrophage accumulation in *Tie2*^+/−^ eyes (Fig. 7D). *Tie2*^+/−^ mice exhibited significantly reduced SC area compared with *Tie2*^+/+^ controls (*Tie2*^+/+^: 1.126 ± 0.074 vs. *Tie2*^+/−^: 0.803 ± 0.038 × 10⁵ μm² per 20× image; unpaired two-tailed Student’s *t* test, *P* = 0.00815) (Fig. 7E). Peri-SC macrophage accumulation was increased in *Tie2*^+/−^ eyes, as reflected by higher IBA1^+^ cell density normalized to SC area (*Tie2*^+/+^: 5.71 ± 0.24 vs. *Tie2*^+/−^: 9.06 ± 0.49 IBA1^+^ cells per 10⁴ μm² SC; unpaired two-tailed Student’s *t* test, *P* = 8.35 × 10⁻⁴) (Fig. 7F). Because *Tie2* haploinsufficiency reproduced the structural and macrophage changes observed with aging, we examined whether it also engaged the macrophage-to-SC *Vegfa* signaling program identified in older eyes. Reference-guided annotation based on the aging dataset identified corresponding Mac1 and Mac5 populations in the 6-week-old *Tie2* scRNA-seq dataset (Supplementary Fig. S7). CellChat analysis predicted Mac1-to-SC *Vegfa*–Vegfr signaling in eyes of young *Tie2*^+/−^ but not *Tie2*^+/+^ mice. However, corresponding Mac5-to-SC *Vegfa–Vegfr* interactions were not detected in either genotype (Fig. 7G). To visualize the expression patterns underlying this interaction-level result, we plotted *Vegfa* expression in Mac1 and Mac5 and *Kdr* expression in SC endothelial cells; these distributions were consistent with the CellChat prediction (Fig. 7H).

### Boosting TIE2 activity with AAV-heptaAng1 improves outflow and delays onset of age-related changes in the SC niche

Because reduced TIE2 signaling produced SC niche changes overlapping with those observed during aging, we asked whether, conversely, boosting TIE2 activity might attenuate age-associated changes. We previously reported that recombinant Hepta-ANGPT1, a soluble ANGPT1 mimetic engineered to robustly activate its cognate receptor, TIE2 (35), transiently lowered IOP in normotensive young adult mice (24). In paired ex vivo perfusion experiments, acute Hepta-ANGPT1 treatment also increased outflow facility by 27% relative to contralateral control eyes (95% CI, 12–44%; P = 0.0042; n = 8; Supplementary Fig. S8), confirming that the IOP-lowering effect was mediated by the conventional outflow pathway. To achieve sustained TIE2 activation for long-term IOP lowering, we generated an AAV vector using scAAV2, which targets the TM and SC inner wall (34, 36) to drive expression of an HA-tagged Hepta-ANGPT1 under control of the constitutively active CMV promoter (hereinafter referred to as AAV-A1, Supplementary Fig. S9A). HEK293 cells showed HA-tag staining 3 days after transduction with AAV-A1, whereas no HA expression was seen in cells transduced with AAV-GFP, confirming protein production (Supplementary Fig. S9B). When delivered to wild-type mice by intracameral injection, AAV-A1 increased p-TIE2 immunostaining intensity in the SC one month after transduction (Supplementary Fig. S9C,D), demonstrating functional Hepta-ANGPT1 activity in the outflow tissues in vivo. We then tested whether this upstream activation improved AHO physiology and attenuated the macrophage/*Vegfa* response in older eyes. Fourteen-month-old wild-type mice received intracameral AAV-A1 in one eye and AAV-GFP in the contralateral eye and were analyzed one month later. IBA1/tracer imaging and quantification, with the SC region delineated based on PECAM1 staining, showed reduced peri-SC macrophage accumulation in AAV-A1-treated eyes, reflected by fewer IBA1^+^ cells per 10^4^ µm^2^ SC (Fig. 8A,B). Following intracameral injection, AAV-A1 increased outflow facility (C_r_; Fig. 8C) and reduced IOP (Fig. 8D) in comparison with matched AAV-GFP controls. *Vegfa* RNAscope confirmed that the proportion of *Vegfa*-expressing IBA1^+^ cells among peri-SC IBA1^+^ macrophages was also reduced in AAV-A1-treated eyes (Fig. 8E,F).

**Figure 8.**
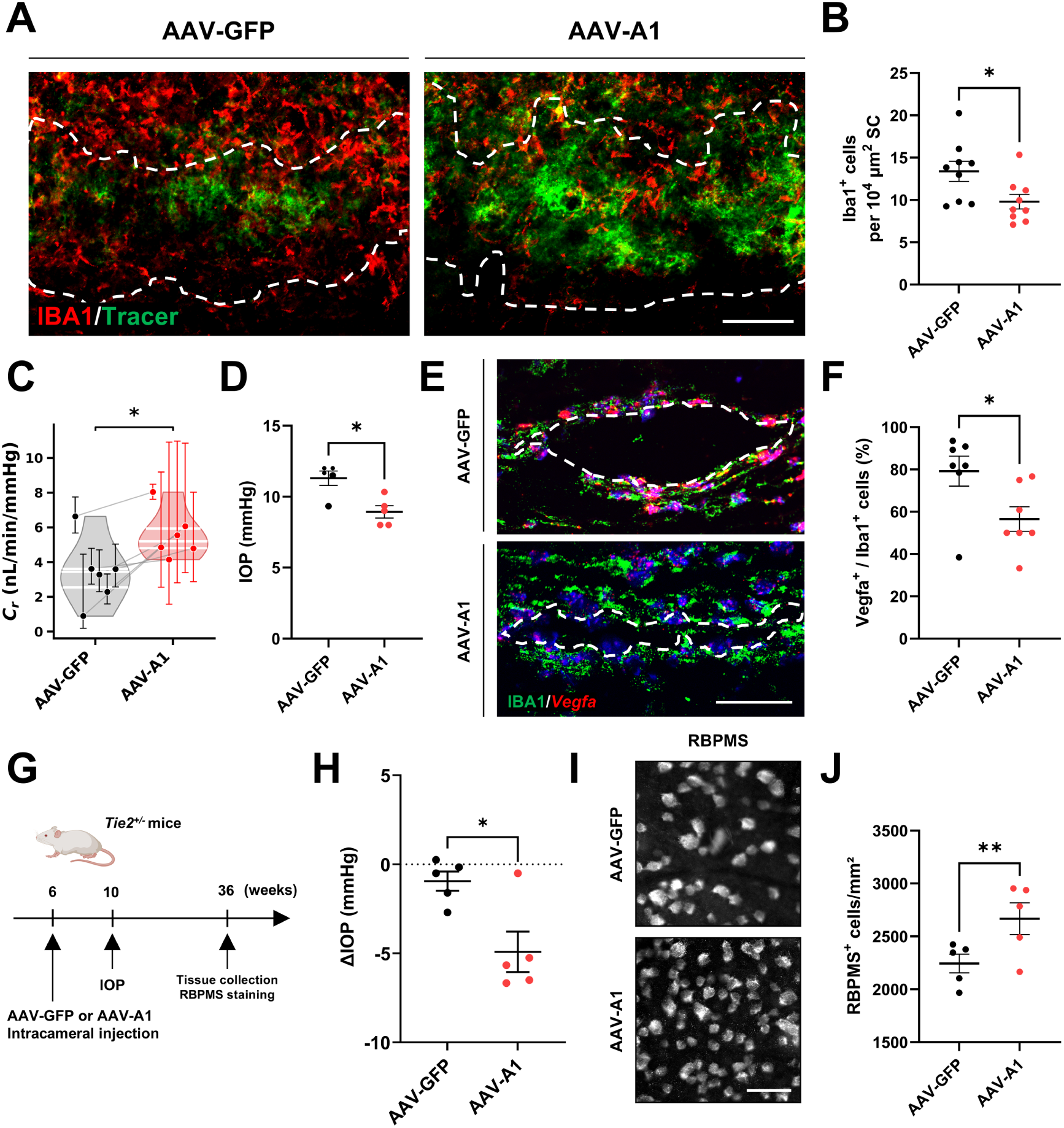
AAV-heptaAng1 improves outflow and reduces macrophage *Vegfa* responses in older eyes and mitigates ocular hypertension and retinal ganglion cell loss in *Tie2*-haploinsufficient mice. (A) Representative IBA1 immunostaining (red) and fluorescent tracer signal (green) in the Schlemm’ s canal (SC) region from older mice treated with AAV-GFP or AAV-A1. Dashed lines delineate the SC based on PECAM1 immunostaining. Scale bar, 100 μm. (B) Quantification of peri-SC IBA1^+^ macrophage density normalized to SC area. (C) Quantification of outflow facility (C_r_) in paired AAV-GFP- and AAV-A1-treated eyes. Raw C_r_ values are shown; statistical analysis was performed using a two-tailed paired t-test on ln(C_r_). (D) Intraocular pressure (IOP) in AAV-GFP- and AAV-A1-treated eyes. (E) Representative *Vegfa* RNAscope (red) combined with IBA1 immunostaining (green) and DAPI counterstaining (blue) in the SC region from AAV-GFP- and AAV-A1-treated eyes. Dashed lines outline the SC region. Scale bar, 50 μm. (F) Quantification of the proportion of *Vegfa*-expressing IBA1^+^ cells among peri-SC IBA1^+^ macrophages around the SC. Data are presented as mean ± SEM unless otherwise indicated. Each dot represents one eye, with AAV-GFP and AAV-A1 measurements paired within each mouse (9 pairs in B, 6 in C, 5 in D, and 7 in F). For (B), (D), and (F), two-tailed paired Student’s t-tests were used (P = 0.0467, 0.0189, and 0.0266, respectively). *P < 0.05, **P < 0.01. (G) Experimental scheme showing intracameral injection of AAV-GFP or AAV-A1 into 6-week-old *Tie2*^+/−^ mice, followed by intraocular pressure (IOP) measurement at 10 weeks of age and retinal ganglion cell analysis at 36 weeks of age. (H) Change in IOP (ΔIOP) at 10 weeks of age relative to baseline in paired AAV-GFP-and AAV-A1-treated *Tie2*^+/−^ mice. (I and J) Representative RBPMS immunostaining images (I) and quantification of RBPMS^+^ retinal ganglion cell density (J) in AAV-GFP-and AAV-A1-treated *Tie2*^+/−^ mice at 36 weeks of age. Scale bar, 50 μm. Data are presented as mean ± SEM. In (H), each dot represents one eye and statistical analysis was performed using a two-tailed paired t-test. In (J), each dot represents one eye (n = 5 pairs) and statistical analysis was performed using a two-tailed paired t-test. *P < 0.05, **P < 0.01.

### TM-targeted ANGPT1 gene therapy mitigated age-related and glaucomatous changes in Tie2-haploinsufficient mice

Because aging alone induced inflammatory remodeling of the SC niche with reduced SC area but without robust ocular hypertension, whereas *Tie2* haploinsufficiency shared key SC phenotypes with aging and produced clear age-dependent IOP elevation, the *Tie2*-haploinsufficient model provided a setting to test whether early TIE2 augmentation could mitigate age-progressive ocular hypertension and retinal ganglion cell (RGC) loss. Consistent with our previous report (33), 9-month-old *Tie2*^+/−^ mice showed reduced peripheral RBPMS^+^ RGC density compared with *Tie2*^+/+^ littermates, whereas central and middle retinal regions were not significantly affected (Supplementary Fig. S10). For this intervention, 6-week-old *Tie2*^+/−^ mice received intracameral AAV-GFP or AAV-heptaAng1 (AAV-A1), with IOP measured at 10 weeks of age and RBPMS staining performed at 36 weeks of age (Fig. 8G). At 10 weeks, the AAV-A1 group showed a significantly greater reduction in IOP than the AAV-GFP group (ΔIOP: −4.92 ± 1.13 vs −0.94 ± 0.54 mmHg, mean ± SEM; two-tailed paired t-test, P = 0.0141; Fig. 8H). RBPMS^+^ retinal ganglion cell density was subsequently quantified at 36 weeks of age to evaluate later retinal preservation. AAV-A1-treated eyes showed a significantly higher density of RBPMS^+^ cells than AAV-GFP-treated controls (2667 ± 150 vs 2244 ± 89 cells/mm^2^, mean ± SEM; two-tailed paired t-test, P = 0.00328; Fig. 8I,J).

## Discussion

### Aging triggers adaptive responses in the SC niche to preserve IOP homeostasis

Older age is a major risk factor for primary open-angle glaucoma (POAG) (3, 4), and prior studies have shown that aging is accompanied by structural alterations in both SC and the trabecular meshwork (TM), including reduced SC dimensions, loss of SC endothelial cells and giant vacuoles, decreased TM cellularity, accumulation of extracellular material, and increased tissue stiffness that are predicted to increase outflow resistance (9, 10, 13–15). Yet in most individuals, these structural and biomechanical changes do not result in elevated IOP or reduced AH outflow facility (9). This apparent dissociation suggests that the aging AH outflow pathway does not simply fail but instead enters a compensated state in which local adaptive mechanisms preserve IOP despite increasing tissue dysfunction. This leaves the tissue vulnerable to additional insults, and ocular hypertension is predicted to emerge when this compensatory capacity becomes insufficient or is compounded by additional genetic, mechanical, or environmental stressors. Our findings identified peri-SC macrophage-derived VEGFA as a compensatory response and showed that increased availability of the TIE2 ligand, ANGPT1, is a strategy to boost SC resilience, delaying the need for VEGFA^+^ macrophage support.

### CX3CR1^+^*Vegfa*/VEGFA^+^ macrophages serve as a niche-specific compensatory program

In other organs, CX3CR1^+^ macrophages have been shown to reside in vascular niches and have been linked to tissue repair (37–40). In the eyes of young mice, resident macrophages located in the trabecular meshwork were shown to be involved in IOP homeostasis, supporting a model whereby tissue-associated macrophages can modulate AH outflow dynamics (20). A companion single-cell study of aging outflow tissues similarly identified increased macrophage abundance and predicted macrophage-derived *Vegfa* signaling to TM cells and inner-wall SC endothelial cells (Balasubramanian et al., concurrent submission). Here, we identified a population of VEGFA-expressing Cx3cr1^+^ macrophages that expands with advancing age adjacent to the SC in mice and is also increased in Tie2-haploinsufficient mice; notably, VEGFA-expressing peri-SC macrophages were also observed in humans. VEGFA is a potent angiogenic and vasoactive factor and has been shown to directly increase outflow facility and reduce SC endothelial barrier resistance through activation of its cognate tyrosine kinase (TK) receptor, VEGFR2, which is expressed in SC endothelial cells (27, 41). Conversely, repeated intravitreal anti-VEGF injections for neovascular retinal diseases have been associated with IOP elevation and reduced aqueous outflow facility (42, 43). VEGFA–VEGFR2 signaling activates eNOS and NO production in endothelial cells, while eNOS-derived NO in SC and downstream endothelial cells promotes conventional outflow facility (44, 45). These data led us to posit that Vegfa-expressing peri-SC macrophages counterbalance age-related structural changes that increase outflow resistance by activating VEGFR2 in SC, promoting vasodilation and reducing barrier resistance. Deletion of *Vegfa* in CX3CR1^+^ cells impaired pressure homeostasis, increasing IOP and reducing outflow facility, consistent with this model.

### Convergence of Adaptive Responses in the Setting of Increased AHO Resistance

In addition to their role in the compensatory response to aging, the observation that Vegfa^+^ macrophages also accumulate in a genetic model of SC dysfunction caused by Tie2 haploinsufficiency was surprising and suggested one of two possibilities: (1) multiple factors that cause vascular stress like SC narrowing and increase outflow resistance trigger a common adaptive response; or (2) reduced TIE2 signaling, which has been linked to vascular aging in other organs (46, 47), is also a driver of age-related changes in the eye and specifically the SC niche.

Although this study cannot distinguish these two possibilities, TIE2 and p-TIE2 have been reported to decline in older SC, whereas pharmacologic TIE2 activation acutely increased SC endothelial proliferation and giant vacuole formation in older mice (48). More broadly, TIE2 supports endothelial quiescence, and reduced TIE2 signaling can promote endothelial activation and immune cell accumulation in other tissues (49).

### Translational implications of boosting TIE2 signaling in the SC

Because our data suggested that POAG occurs when normal compensatory mechanisms become insufficient to maintain healthy IOP, we sought to identify an SC-targeted therapeutic approach to promote resilience. ANGPT1/TIE2 signaling is required for SC development and adult maintenance (48, 50), and TIE2 preserves endothelial quiescence in other vascular beds (49). While macrophage-derived Vegfa was found to be protective, direct VEGFA delivery to the eye is problematic due to the narrow safety window and risk of neovascular glaucoma (51–54). Instead, because reduced TIE2 signaling was associated with SC remodeling that shared features with age-associated changes and because levels of TIE2 and p-TIE2 within the canal are known to decline with age (48), we wondered if boosting TIE2 activity would attenuate age-associated phenotypes in wild-type mice, thereby extending resilience even in the setting of additional ‘hits’ to the tissues of the anterior chamber. We have previously reported that intravitreal injection of a potent TIE2 ligand mimetic – heptaAng1 can lower IOP transiently in adult wild-type mice (24). However, to be effective clinically, a more sustained effect is needed. To accomplish this, we developed a gene therapy approach, using an AAV that transduces cells in the TM, the physiologic location of endogenous angiopoietin production (36, 50). In older eyes, AAV-heptaAng1 increased outflow facility, lowered IOP, and reduced peri-SC macrophage density and specifically reduced the proportion of *Vegfa*^+^ cells among these macrophages, supporting the model that elevated TIE2 signaling relieves vascular stress that triggers this compensatory response. Furthermore, in *Tie2*-haploinsufficient mice, early AAV-heptaAng1 treatment was associated with long-lasting preservation of RGC density, supporting its translational potential for sustained benefit. While future studies in other glaucoma models and human tissues are needed to determine whether TIE2 signaling deficiency (reduced p-TIE2 in SC) itself drives age-related changes or if TIE2 activity serves as a general ‘vasculoprotective’ factor (49), these findings support a beneficial role for TIE2 activation, consistent with the strong genetic data linking this pathway to risk of both developmental (primary congenital glaucoma) and adult-onset glaucomas (POAG) (55–57).

### Study limitations

While our study identified a clear role for Vegfa^+^ macrophages in maintenance of the aging outflow pathway, several limitations should be considered. First, *Cx3cr1*-CreERT2 targets CX3CR1-lineage cells rather than macrophages exclusively, and we did not directly define the ontogeny, turnover, or long-term fate of the CX3CR1^+^*Vegfa/VEGFA*^+^ population, although the long-lasting effect of Vegfa deletion following a short course of tamoxifen supports a contribution from long-lived CX3CR1-lineage cells to the adaptive response. Second, although CX3CR1-lineage *Vegfa deletion* supports an outflow-protective role for macrophage-associated VEGFA, the downstream signaling pathways that mediate this effect remain incompletely defined. Finally, the mechanisms by which aging reduces TIE2 signaling and induces the macrophage *Vegfa response* remain unknown.

## Conclusion

In sum, these findings support a model (Fig. 9) whereby peri-SC *Vegfa*^+^ macrophages serve as one compensatory mechanism of age-related outflow adaptation and demonstrate that boosting TIE2 activity is an exciting therapeutic strategy to promote SC resilience, delaying the need for compensatory adaptation.

**Figure 9.**
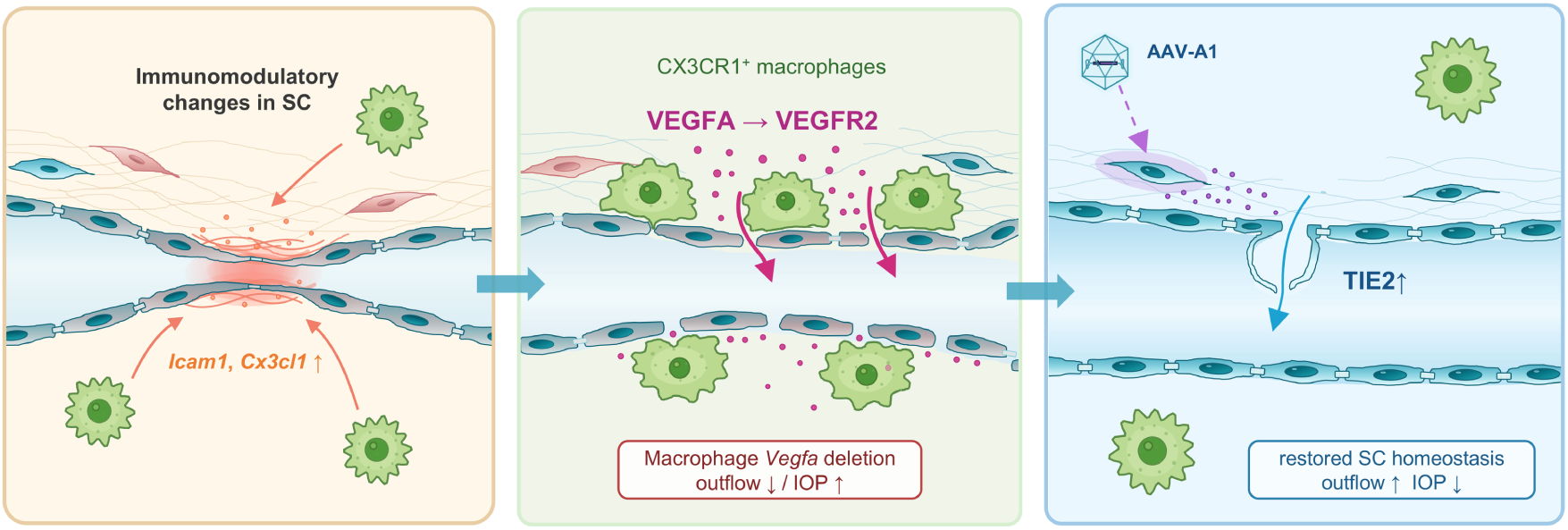
Proposed model of aging- and TIE2-dependent remodeling of the Schlemm’s canal niche. Aging remodels the SC niche with reduced SC area, immunomodulatory endothelial changes, and accumulation of peri-SC CX3CR1^+^ macrophages, while reduced TIE2 signaling converges on reduced SC area and peri-SC macrophage accumulation. Macrophage-derived VEGFA appears to act as a compensatory response that partially supports outflow, whereas AAV-heptaAng1 treatment restores SC homeostasis, reduces peri-SC macrophages and *Vegfa*-expressing macrophages, improves outflow, lowers IOP, and preserves RGCs.

## Materials and Methods

### Sex as a biological variable

Both male and female mice were used unless otherwise indicated. Analyses were not stratified by sex. Sex information was not available for the human donor tissues.

### Animal generation and husbandry

All mice were housed in the Center for Comparative Medicine at Northwestern University (Chicago, IL, USA) under standard conditions (12-hour light/dark cycle, ambient temperature 21–23 °C, 30–70% relative humidity) with ad libitum access to water and standard chow (Teklad #7912, Envigo, Indianapolis, IN, USA). Wild-type C57BL/6J mice were used for aging studies unless otherwise indicated. For *Tie2* haploinsufficiency experiments, *Tie2* heterozygous-null mice (*Tie2*^+/−^) were generated by EIIa-Cre-mediated germline recombination. Briefly, *Tie2* floxed mice (*Tie2*^fl/fl^), previously described by Savant et al. (58) and generated by Ingenious Targeting Laboratories, were crossed with EIIa-Cre mice (Tg(EIIa-cre)C5379Lmgd/J, JAX #003724) to generate germline-deleted *Tie2* alleles, and the Cre transgene was subsequently removed by crossing to wild-type mice. *Tie2*^+/−^ mice were then maintained and compared with *Tie2*^+/+^ littermate controls. Genotyping was performed by PCR using the following primers: *Tie2* forward, 5′-TTTCCATCACACGGGCTTTG-3′; *Tie2* reverse, 5′-TCAAAACCGTTGCCATGTGT-3′; Cre forward, 5′-GTGCAAGTTGAATAACCGGAAATGG-3′; and Cre reverse, 5′-AGAGTCATCCTTAGCGCCGTAAATCAAT-3′.

For macrophage lineage-labeling experiments, eyes from *Cx3cr1*-CreERT2; ZsGreen reporter mice of different ages were obtained from the Budinger laboratory. This tamoxifen-inducible reporter system has been described previously (59). Briefly, *Cx3cr1*-CreERT2 mice (JAX #020940) were crossed with the ZsGreen reporter line Ai6 (JAX #007906). Tamoxifen was administered by oral gavage (10 mg/100 µL) twice, 7 and 2 days before tissue harvest, to induce reporter expression.

For CX3CR1^+^ macrophage-targeted *Vegfa* deletion experiments, we newly generated a conditional *Vegfa* allele in this study. With assistance from the Northwestern University Transgenic and Targeted Mutagenesis Laboratory, we designed a CRISPR-Cas9 strategy to target intron 2 and intron 5 of the *Vegfa* locus and introduce two loxP sites (Supplementary Fig. S5A). Specifically, one CRISPR RNA (crRNA) was designed for each target site, together with 190-mer and 212-mer single-stranded oligodeoxynucleotide donor templates (ssODNs) containing the loxP sequence and an adjacent XmaI recognition site; these sequences are provided in Supplementary Table S3. crRNA and tracrRNA were annealed at 95 °C for 3 min to generate guide RNA (gRNA), and within 5 h after in vitro fertilization, one-cell-stage embryos obtained from 6–8-week-old C57BL/6N females (Charles River) were introduced into an electroporation solution containing 100 ng/µL gRNA, 20 ng/µL ssODN, and 100 ng/µL HiFi Cas9 in nuclease-free duplex buffer. CRISPR electroporation was performed using an ECM 830 Square Wave Electroporation System, and only embryos that developed to the 2-cell stage the following day were transferred into the oviducts of pseudopregnant females. Founder mice were screened by Sanger sequencing of the modified region to confirm the intended loxP insertions (Supplementary Fig. S5B). Genotyping of the *Vegfa* floxed allele was performed using tail DNA and PCR with *Vegfa* F (5′-CCGTCTCTGGAATATGGGCA-3′) and *Vegfa* R (5′-CGTCCATCCCCTGATACACA-3′), which yielded a 552-bp product for the wild-type allele and a 587-bp product for the floxed allele. When needed, BglI digestion was used for confirmation, yielding 305-bp and 247-bp fragments for the wild-type allele and an undigested 587-bp fragment for the floxed allele (Supplementary Fig. S5C). These *Vegfa*^fl/fl^ mice were then crossed with *Cx3cr1*-CreERT2 knock-in mice (B6.129P2(Cg)-*Cx3cr1*tm2.1(cre/ERT2)Litt/WganJ, JAX #021160) to generate *Cx3cr1*-CreERT2; *Vegfa*^fl/fl^ conditional knockout mice. To induce recombination, tamoxifen (75 mg/kg/day) was administered intraperitoneally for 3 consecutive days starting at 6 weeks of age. Both male and female mice were used unless otherwise indicated.

### Single-cell RNA-seq

Single-cell RNA-seq of murine limbal/iridocorneal angle tissues was performed broadly according to our previously published workflow for outflow tissues (23), with modifications for the present study. Briefly, limbal/iridocorneal angle tissues containing SC, TM, and adjacent tissues were microdissected from enucleated eyes and enzymatically dissociated into single-cell suspensions as previously described (23). For the aging dataset, tissues from young (6-week-old) and older (15-month-old) mice were analyzed, with six eyes pooled per sample to generate two independent biological samples per group. For the *Tie2* dataset, limbal/iridocorneal angle tissues from *Tie2*^+/−^ and control mice were processed in an analogous manner, with six eyes pooled per sample to generate two independent biological samples per group. Single-cell suspensions were submitted to the Northwestern University NUSeq Core Facility for library preparation using the 10x Genomics Chromium Next GEM Single Cell 3′ Reagent Kits v3.1 and sequenced on an Illumina HiSeq 4000 instrument. Raw sequencing data were processed using Cell Ranger (version 7.1.0, 10x Genomics) with alignment to the mouse reference genome (mm10). For the aging dataset, cells with fewer than 200 or more than 6,000 detected features or with mitochondrial transcript content greater than 10% were excluded. Doublets were identified using scDblFinder (version 1.20.0; nfeatures = 1000), and only singlets were retained. Each sample was normalized using NormalizeData, and 2,000 variable features were identified using the vst method. Datasets were integrated using SelectIntegrationFeatures, FindIntegrationAnchors, and IntegrateData. For the aging dataset, the integrated assay was scaled and subjected to principal component analysis using 50 PCs, followed by JackStraw analysis, UMAP, graph-based neighbor finding using dims = 1:50, and clustering at a resolution of 0.6. For the *Tie2* dataset, the integrated assay was subjected to principal component analysis using 50 PCs, followed by UMAP, graph-based neighbor finding using dims = 1:30, and clustering at a resolution of 0.5. Downstream analyses were performed in Seurat (version 5.4.0). Cell identities were assigned based on established marker genes and our prior iridocorneal angle single-cell atlas (24). For aging analyses, the SC endothelial cluster was identified and subsetted for differential expression analysis between young and older SC cells using FindMarkers with the Wilcoxon rank-sum test, with a log fold change threshold of 0.25 and an adjusted P value threshold of 0.05. Enrichr pathway analysis was performed using the top 200 genes upregulated in older SC cells. For myeloid analyses, monocytes and macrophages were subsetted and reclustered to resolve Mono and Mac1–Mac5 subpopulations. In the *Tie2* dataset, myeloid cells were annotated by reference-guided transfer using the aging dataset as reference. Cell-cell communication analysis was performed using CellChat (version 1.6.1) across all annotated cell types. Interactions involving SC and myeloid populations were subsequently extracted for visualization, and interaction strength was evaluated using the interaction weight metric.

### Outflow facility measurement

Conventional outflow facility was measured ex vivo in freshly enucleated mouse eyes using a constant-pressure perfusion system (iPerfusion 3), broadly as previously described (60, 61). Eyes were cannulated through the anterior chamber with a glass needle under a stereomicroscope and submerged in temperature-controlled PBS supplemented with 5.5 mM glucose. After cannulation, eyes were held at approximately 8 mmHg for a 30–45 min acclimation period and then perfused using a multistep pressure protocol spanning the physiological range. Flow rate (Q) and pressure (P) were recorded continuously, and steady-state values at each pressure step were identified programmatically to generate one Q–P data point per step. The flow-pressure relationship was fit using the power-law model Q(P) = C_r_ · P · (P/P_r_)^β^, with P_r_ fixed at 8 mmHg, where C_r_ represents the reference outflow facility (nL/min/mmHg). For statistical comparisons, C_r_ values were log-transformed before analysis. For acute Hepta-ANGPT1 perfusion experiments, eyes from male C57BL/6J mice (11 to 12.5 weeks old) were analyzed in a paired design, with one eye receiving 1 μg/mL Hepta-ANGPT1 in Dulbecco’s phosphate-buffered saline with calcium and magnesium, supplemented with 5.5 mM D-glucose (DBG) and the contralateral eye receiving DBG alone. Both eyes were perfused simultaneously. Following 20 min at 8 mmHg, perfusate was exchanged at 5 μL/min for 100 μL, as previously described (62), followed by an additional 30 min at 8 mmHg before initiation of the pressure-step protocol. Assignment of control and treated eyes to the two perfusion systems was alternated across pairs to minimize system-dependent bias. Relative differences in C_r_ were calculated between treated and contralateral control eyes and analyzed using the weighted paired t-test implemented in iPerfusion, which accounts for uncertainty in the fitted Cr estimate for each eye (60).

### IOP measurements

IOP measurements were performed in awake mice between 9 and 11 AM using an iCare TonoLab rebound tonometer as previously described (24). Cohorts of mutant mice with littermate controls were measured at each reported time point. Unless otherwise indicated in the figure legends, IOP values from left and right eyes were averaged to obtain values reported in the manuscript.

### Immunofluorescence staining

Mouse anterior segment flat mounts and retinal flat mounts were processed broadly as previously described (50, 63, 64). Briefly, eyes were enucleated and immersion-fixed in 2% paraformaldehyde overnight at 4 °C. For anterior segment flat mounts, eyes were bisected from the optic nerve to the center of the cornea, and the lens and retina were removed. Tissues were blocked in TBS containing 5% donkey serum, 2.5% bovine serum albumin (BSA), and 0.5% Triton X-100 overnight at 4 °C, followed by overnight incubation at 4 °C with primary antibodies diluted in the same buffer. After washing, tissues were incubated with species-appropriate secondary antibodies diluted in blocking buffer, counterstained with DAPI, flat-mounted, and imaged using a Nikon A1R confocal microscope. Image processing and quantitative analyses were performed using Fiji software (ImageJ 1.54p; Java 1.8.0_322, 64-bit). For human analyses, de-identified residual limbal/corneal rim tissues containing SC and TM were obtained after the central cornea was used for deceased-donor transplantation. Cryosections were prepared for PECAM1/IBA1 immunostaining, and donor age and death-to-preservation interval are summarized in Supplementary Table S1. Antibody information is provided in Supplementary Table S4.

### RNAscope in situ hybridization

*Vegfa*/*VEGFA* mRNA was detected by fluorescent in situ hybridization using the RNAscope Multiplex Fluorescent Reagent Kit v2 (Advanced Cell Diagnostics [ACD], Bio-Techne; Cat. No. 323100), according to the manufacturer’s instructions. The Mm-*Vegfa* probe (Cat. No. 312931) was used for mouse samples, and the Hs-*VEGFA* probe (Cat. No. 423161) was used for human samples. Sample pretreatment was performed using RNAscope Pretreatment Reagents (Cat. No. 322381) according to the manufacturer’s recommendations for the corresponding sample type. Briefly, sections mounted on SuperFrost Plus slides were subjected to hydrogen peroxide treatment, target retrieval and protease digestion as appropriate, followed by probe hybridization for 2 h at 40 °C, sequential amplification (AMP1–AMP3), HRP-based fluorescent signal development, DAPI counterstaining, and mounting. After RNAscope development, sections were subjected to immunolabeling with anti-CD68 or anti-Iba1 as indicated in each experiment. Antibody and probe information is provided in Supplementary Table S4.

### Intracameral injection

Mice were anesthetized with isoflurane (3% for induction and 1.5–2% for maintenance) and placed on a heated platform. Intracameral injections were performed using a 10 μL NanoFil syringe (World Precision Instruments) fitted with a 35G beveled needle (NF35BV, World Precision Instruments), mounted on an UltraMicroPump (UMP3, World Precision Instruments) connected to a MICRO-2T SMARTouch controller and micromanipulator. For tracer-labeling experiments, FluoSpheres™ (20 nm, carboxylate-modified polystyrene; Thermo Fisher Scientific) were diluted in sterile PBS containing Ca^2+^/Mg^2+^ to a final concentration of 1 × 10^11^ particles/mL, and 1 μL was injected into the anterior chamber. Eyes were collected 30 min after injection and immersion-fixed overnight in 2% paraformaldehyde at 4 °C. For older-eye AAV-A1 experiments, ultra-purified recombinant scAAV2 vectors generated by VectorBuilder Inc. (Chicago, IL, USA) were used: scAAV2-CMV-2×HA-ANGPT1-C4BP (AAV-A1; VB250306-1238kzg) and a matching scAAV2-CMV-EGFP control vector (AAV-GFP; VB010000-9304aud). AAV-GFP or AAV-A1 was injected into the anterior chamber at 1 μL per eye (5 × 10^12^ GC/mL), and eyes were analyzed 1 month later unless otherwise indicated. Viral genome titers were determined by qPCR targeting the AAV2 ITR sequence according to the manufacturer’s certificate of analysis.

### In vitro AAV expression validation

HEK293 cells were used for heterologous expression validation, as in prior ANGPT1 expression assays (50). Cells were seeded in 48-well plates and exposed to control AAV-GFP or AAV-heptaAng1 (AAV-A1). Three days later, cells were analyzed by fluorescence imaging for GFP fluorescence and HA-tag signal.

### SC and RGC immunofluorescence imaging and quantification

For SC analyses, enucleated eyes were fixed overnight in 2% paraformaldehyde (PFA) at 4 °C. After removal of the posterior segment and lens, anterior segments were prepared as flat mounts and immunostained as described above. After staining, additional small relaxing cuts were made around the cornea to facilitate flattening of the SC region, and tissues were mounted with the outer scleral surface facing the coverslip. Confocal z-stacks were acquired from one central field in each quadrant (four fields per eye) using a Nikon A1 confocal microscope with a 20× objective lens. Maximum-intensity projections were generated using Fiji software (ImageJ 1.54p; Java 1.8.0_322, 64-bit), and SC area and peri-SC immune-cell measurements were quantified within the SC region. Depending on the experiment, the CD45^+^ area, numbers of IBA1^+^ cells, *Cx3cr1*^+^ cells, *Vegfa*-expressing CD68^+^ cells, or *Vegfa*-expressing IBA1^+^ cells were normalized to SC area or to circumferential SC length, as indicated in the corresponding figure legends. All acquisition settings were kept identical across groups, and image analysis was performed in a blinded manner. For segmental outflow analysis, maximum-intensity projections were generated in Fiji, and the SC region was manually delineated as the region of interest based on PECAM1 staining. Pearson’s correlation coefficient (r) between the fluorescent tracer and IBA1 channels within the SC region was calculated using the Coloc 2 plugin in Fiji, with the “Pearson’s R value above threshold” output to reduce background signal. Values from four quadrants per eye were averaged across both eyes, yielding one mouse-level value from eight quadrants.

For RGC analyses, retinas were dissected, flat-mounted, and immunostained as described above. Images were acquired using a Nikon Ti2 microscope at 20× magnification from central, middle, and peripheral retinal regions (0.1, 0.8, and 1.5 mm from the optic disc, respectively), with one field per quadrant at each eccentricity (12 fields per retina). Images were cropped to 200 × 200 μm, RBPMS^+^ cells were manually counted, and RGC density was calculated as cells/mm^2^.

## Statistical analysis

Statistical analyses were performed using GraphPad Prism 10.0.4 (GraphPad Software, San Diego, CA, USA) and R version 4.4.2. Data are presented as mean ± SEM unless otherwise indicated. Unless otherwise indicated, two-group comparisons were performed using unpaired two-tailed Student’s t-tests. Paired two-tailed t-tests were used for paired-eye experiments. For comparisons among multiple groups, one-way ANOVA followed by Dunnett’s multiple-comparisons test was used when each group was compared with a single control, and two-way ANOVA followed by Šídák’s multiple-comparisons test was used for experiments involving two independent variables when pairwise comparisons were performed. Pearson correlation analysis was used for correlation analyses. The specific statistical test used for each dataset is indicated in the corresponding figure legend. Because outflow facility (C_r_) is log-normally distributed (65), group comparisons were performed using natural log-transformed values, ln(C_r_), as previously described (60). A two-sided P < 0.05 was considered statistically significant. P values are denoted as follows: *P < 0.05, **P < 0.01, ***P < 0.001, and ****P < 0.0001. All analyses were performed in a blinded fashion with respect to genotype or treatment whenever feasible.

## Study approvals

All animal experiments were approved by the Animal Care and Use Committee at Northwestern University (Evanston, IL, USA) and were performed in accordance with the ARVO Statement for the Use of Animals in Ophthalmic and Vision Research. Human donor limbal/corneal rim tissues used for immunostaining analyses were de-identified residual tissues from deceased donors that remained after corneal transplantation.

## Data Availability

The scRNA-seq data generated in this study have been deposited in the NCBI Gene Expression Omnibus (GEO) under accession number GSE341678. Additional data supporting the findings of this study are available from the corresponding author upon reasonable request. Values for all data points in graphs are provided in the Supporting Data Values file.

## Author Contributions

NK, BRT, and SEQ designed the study. NK, GR, and ERT conducted experiments, and NK, YZ, ERT, and BRT analyzed data. DKD, TO, HLL, CER, RSF, HJL, HG, and GRSB contributed research materials or technical support. DRO, HG, GRSB, BRT, and SEQ contributed to data interpretation. SEQ supervised the study. NK drafted the manuscript, and YZ, ERT, HG, GRSB, BRT, and SEQ contributed to manuscript revision.

## Generative AI program use

OpenAI Codex/ChatGPT (OpenAI; GPT-5) was used on multiple occasions during manuscript preparation and revision, most recently in August 2026, to assist with language editing. The tool was used to improve clarity, concision, and consistency of wording. It was not used to generate scientific hypotheses, design experiments, analyze data, interpret results, create figures, or generate original scientific conclusions. All AI-assisted edits were reviewed, revised as needed, and approved by the authors, who take full responsibility for the manuscript content.

## Acknowledgements

Single-cell RNA sequencing was conducted at the Northwestern University NUSeq Core Facility. Confocal imaging was performed at the Center for Advanced Microscopy of the Feinberg School of Medicine, supported by NCI CCSG P30 CA060553. We thank Jeremy A. Lavine and Edward B. Thorp for helpful discussions and valuable scientific input. We thank Joseph van Batenburg-Sherwood for setting up the iPerfusion outflow facility system used in this study. The mutant mice used in this study were generated by the Northwestern University Transgenic and Targeted Mutagenesis Laboratory (RRID: SCR_017772), which is supported in part by NIH grant CA60553 to the Robert H. Lurie Comprehensive Cancer Center at Northwestern University.

