## Supplementary information for "VEGFA-Positive Macrophages Regulate Aqueous Humor Outflow in Aged Mice and Humans"

### Supplementary Figures

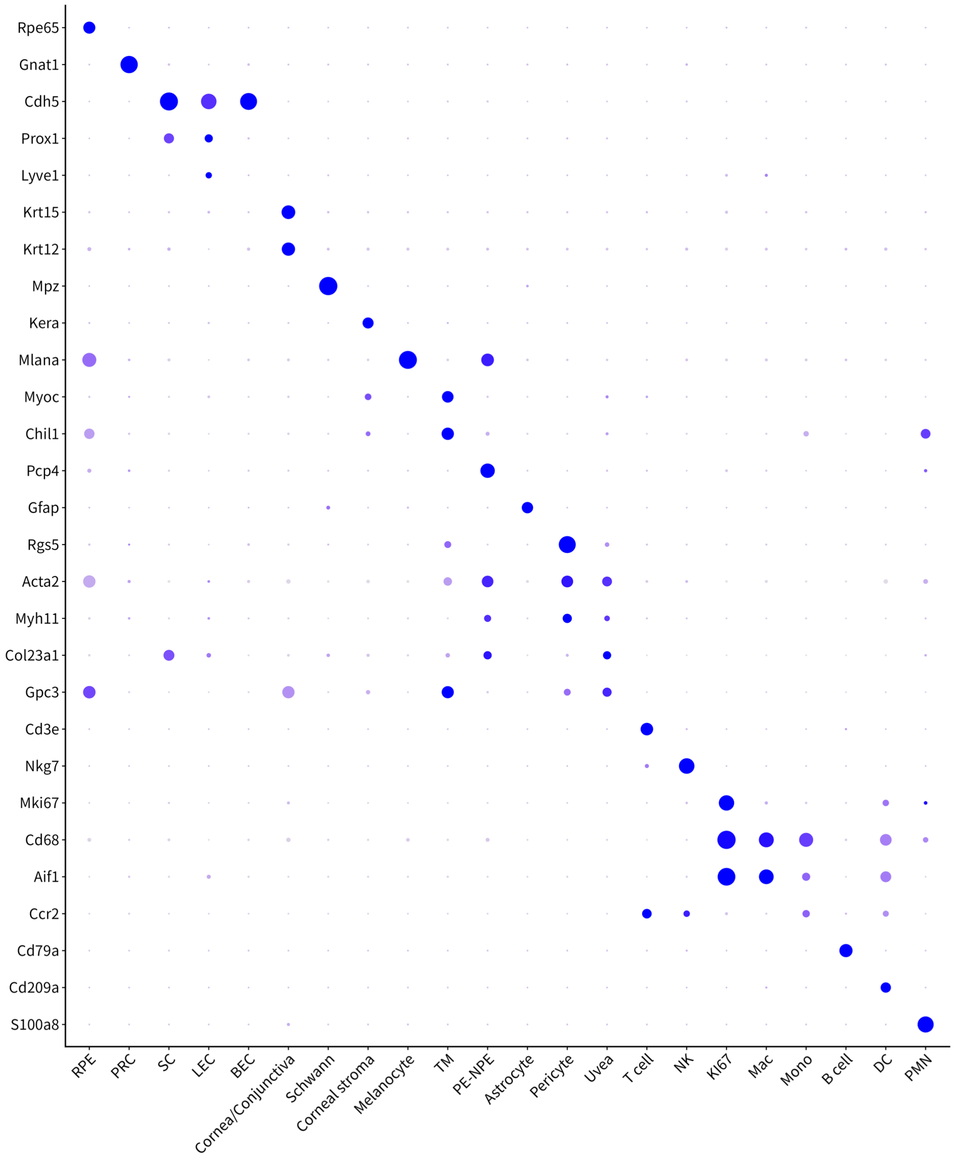


**Supplementary Figure S1. Representative marker genes used for annotation of anterior segment cell populations in the integrated single-cell RNA-seq dataset.**

Dot plot showing expression of selected canonical marker genes across the major cell clusters identified in limbal/iridocorneal angle tissues. These markers support the cell-type annotations shown in Figure 1B. Dot size indicates the percentage of cells expressing each gene, and dot color indicates average expression level.

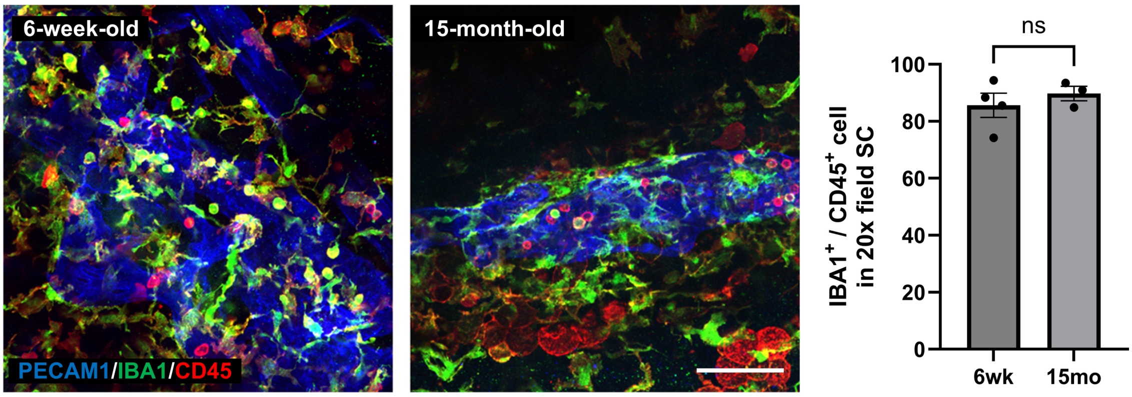


**Supplementary Figure S2. The majority of CD45⁺ leukocytes in the Schlemm's canal region are IBA1⁺ at both young and older time points.**

Representative PECAM1/IBA1/CD45 immunostaining images of the Schlemm's canal (SC) region showing overlap between IBA1 and CD45 signals in peri-SC immune cells. Quantification of the proportion of CD45⁺ cells that were IBA1⁺ in the SC region showed no significant difference between 6-week-old and 15-month-old mice. Scale bar, 100 μm. Data are presented as mean ± SEM. Student's two-tailed unpaired t-test was used. ns, not significant.

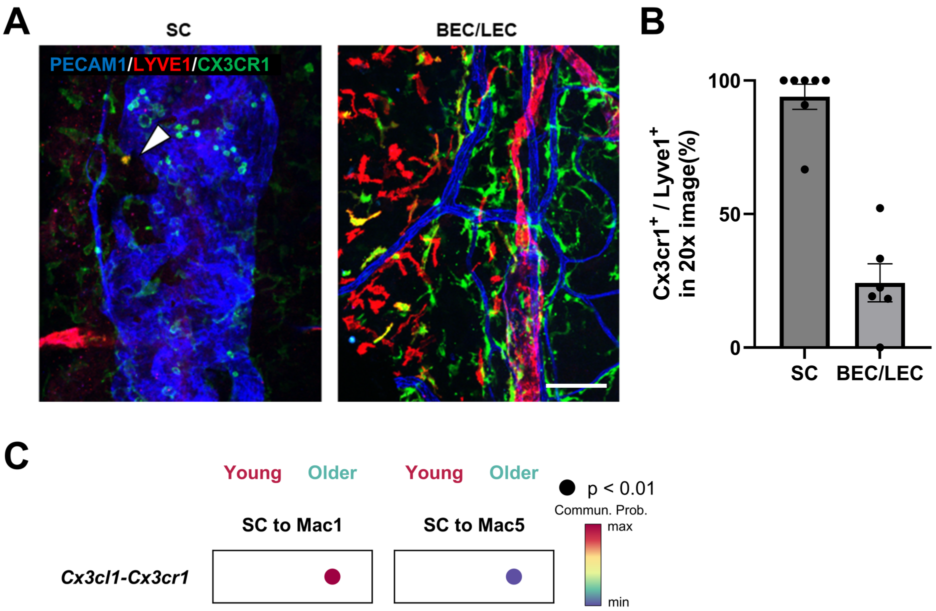


**Supplementary Figure S3. CX3CR1 and LYVE1 expression patterns in macrophages across the conventional outflow pathway.**

(A) Representative PECAM1/LYVE1/CX3CR1 immunostaining images of the Schlemm's canal (SC) region and of blood endothelial cell/lymphatic endothelial cell (BEC/LEC) regions. In the SC region, LYVE1$^{+}$ cells were rare, but when present they were predominantly CX3CR1$^{+}$ (arrowhead). In contrast, LYVE1$^{+}$ cells in the BEC/LEC region showed greater heterogeneity in CX3CR1 expression. Scale bar, 100 μm. (B) Quantification of the proportion of LYVE1$^{+}$ cells that were CX3CR1$^{+}$ in the indicated regions. Each dot represents one image. Statistical analysis was performed using a two-tailed unpaired Student's t-test. $P < 0.0001$. Data are presented as mean ± SEM. (C) Predicted *Cx3cl1*–*Cx3cr1* signaling between Mac1/Mac5 and SC in young versus older eyes (color, probability; black circle, $P < 0.01$).

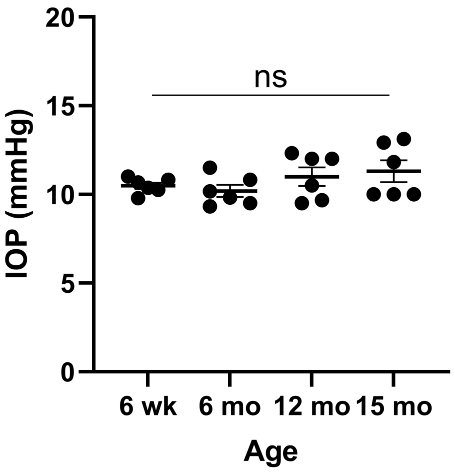


**Supplementary Figure S4. Intraocular pressure does not show a significant age-dependent change in mice.**

Intraocular pressure (IOP) was measured in mice at 6 weeks, 6 months, 12 months, and 15 months of age. No significant difference was detected among age groups. Data are presented as mean ± SEM. One-way ANOVA was used. ns, not significant.

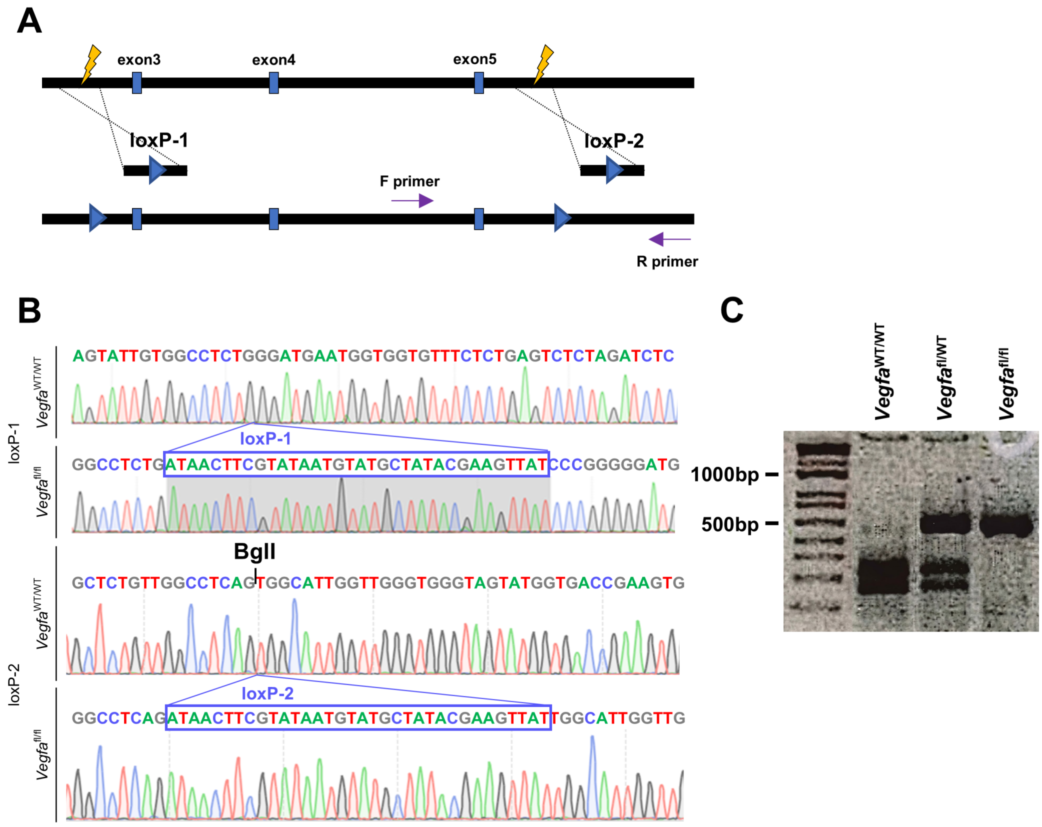


**Supplementary Figure S5. Generation and validation of the *Vegfa* floxed allele.**

(A) Schematic of the CRISPR-Cas9 targeting strategy used to introduce loxP sites into intron 2 (loxP-1) and intron 5 (loxP-2) of the *Vegfa* locus, thereby flanking exons 3–5. Purple arrows indicate the primers used for genotyping PCR. (B) Representative Sanger sequencing chromatograms confirming correct insertion of loxP-1 and loxP-2 in the *Vegfa* floxed allele. The inserted loxP sequences are boxed. (C) Representative PCR/BglI-based genotyping of *Vegfa* alleles. After BglI digestion, the wild-type allele yielded 305-bp and

247-bp fragments, whereas the floxed allele remained undigested at 587 bp; heterozygous mice showed both patterns. Lane order is *Vegfa*$^{wt/wt}$, *Vegfa*$^{wt/fl}$, and *Vegfa*$^{fl/fl}$.

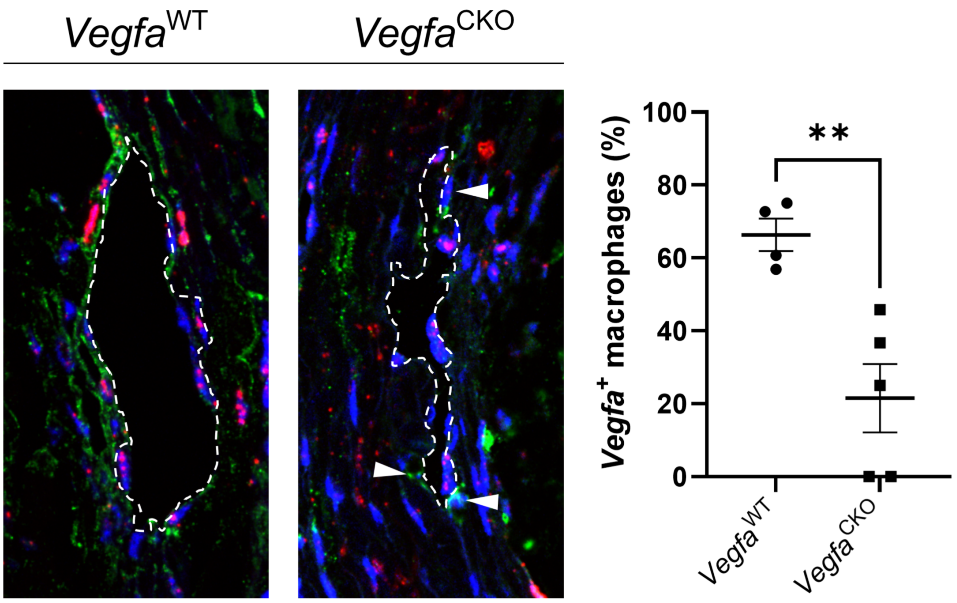


**Supplementary Figure S6. *Vegfa* expression is reduced in Cx3cr1⁺ macrophages in 9-month-old *Vegfa* CKO eyes.**

Representative *Vegfa* RNAscope (red) combined with IBA1 immunostaining (green) and DAPI (blue) images of the Schlemm's canal (SC) region from *Vegfa* WT and *Vegfa* CKO mice harvested at 9 months of age. Dashed lines outline the SC region. Arrowheads indicate peri-SC IBA1⁺ macrophages. Quantification shows the percentage of *Vegfa*-expressing macrophages among peri-SC IBA1⁺ macrophages around the SC. Data are presented as mean ± SEM. Each dot represents a mouse-level value generated by averaging image-level measurements where applicable (WT n = 4, CKO n = 5). Student's two-tailed unpaired t-test was used. **P < 0.01.

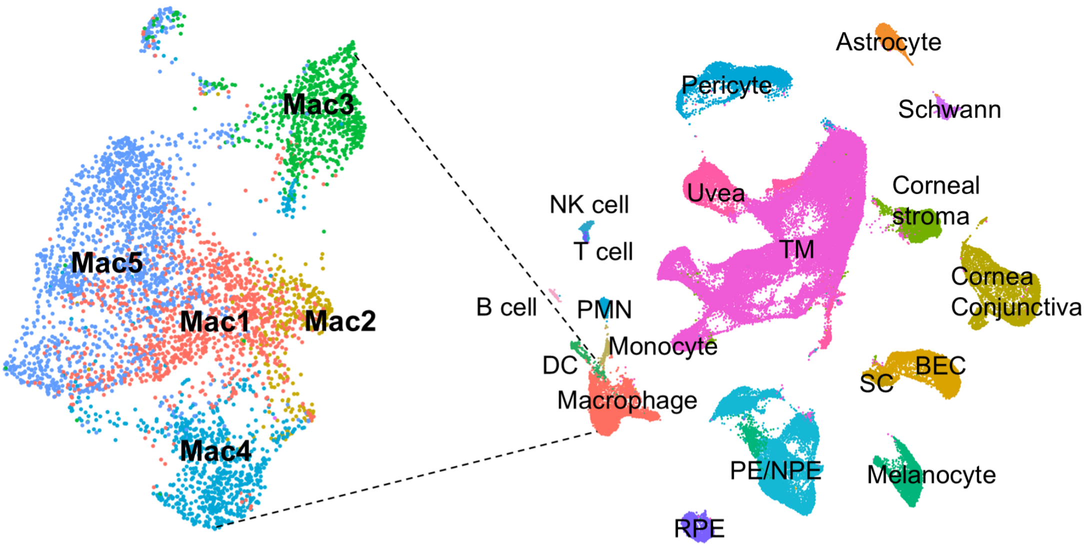


**Supplementary Figure S7. Reference-guided annotation of myeloid populations in the *Tie2* dataset identifies monocytes and five macrophage subclusters.**

UMAP of the integrated *Tie2* anterior segment dataset showing major cell populations (right) and magnified view of the myeloid compartment (left). Reference-guided transfer annotation based on the aging dataset resolved monocytes (Mono) and five macrophage subclusters (Mac1–Mac5), enabling cross-dataset comparison of myeloid cell states. Dashed lines indicate the myeloid compartment selected for magnified visualization.

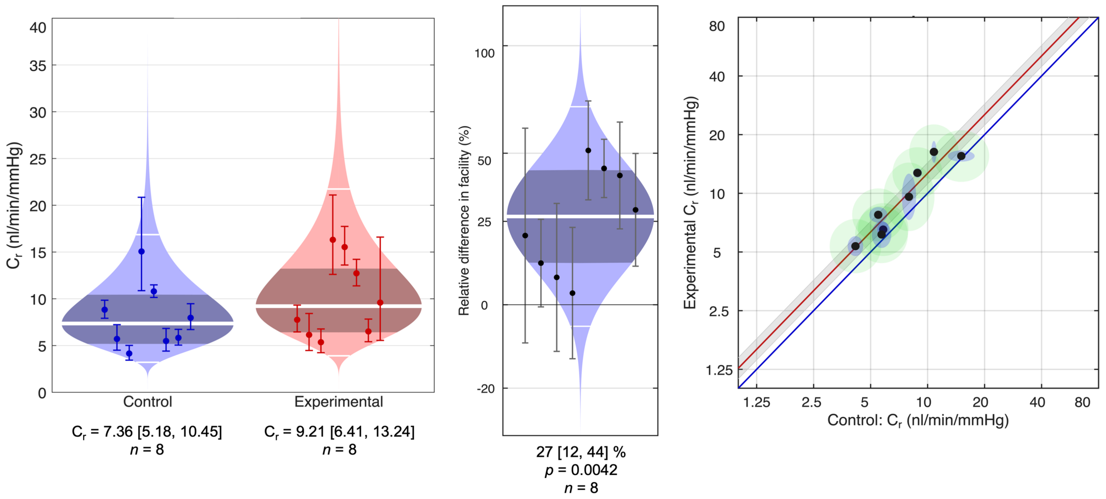


**Supplementary Figure S8. Acute Hepta-ANGPT1 treatment increases conventional outflow facility in paired ex vivo eye perfusion.**

Eyes from male C57BL/6J mice were analyzed in a paired design, with one eye perfused with Hepta-ANGPT1 and the contralateral eye perfused with Dulbecco's phosphate-buffered saline with calcium and magnesium, supplemented with 5.5 mM D-glucose (DBG) alone. Left, absolute conventional outflow facility (Cr) values for control and Hepta-ANGPT1-treated eyes. White lines indicate geometric means and dark shaded areas indicate 95% confidence intervals. Middle, paired relative difference in facility, showing a 27% increase in treated eyes relative to contralateral controls (95% CI, 12–44%; P = 0.0042; n = 8). Right, scatter plot of treated versus control Cr values. The red line indicates the fitted relationship with its 95% confidence interval (gray), and the blue line indicates the line of identity. All Hepta-ANGPT1-treated eyes exhibited higher Cr than their paired control eyes. Statistical analysis of the paired relative difference was performed using the weighted paired t-test implemented in iPerfusion, which accounts for uncertainty in the fitted Cr estimate for each eye.

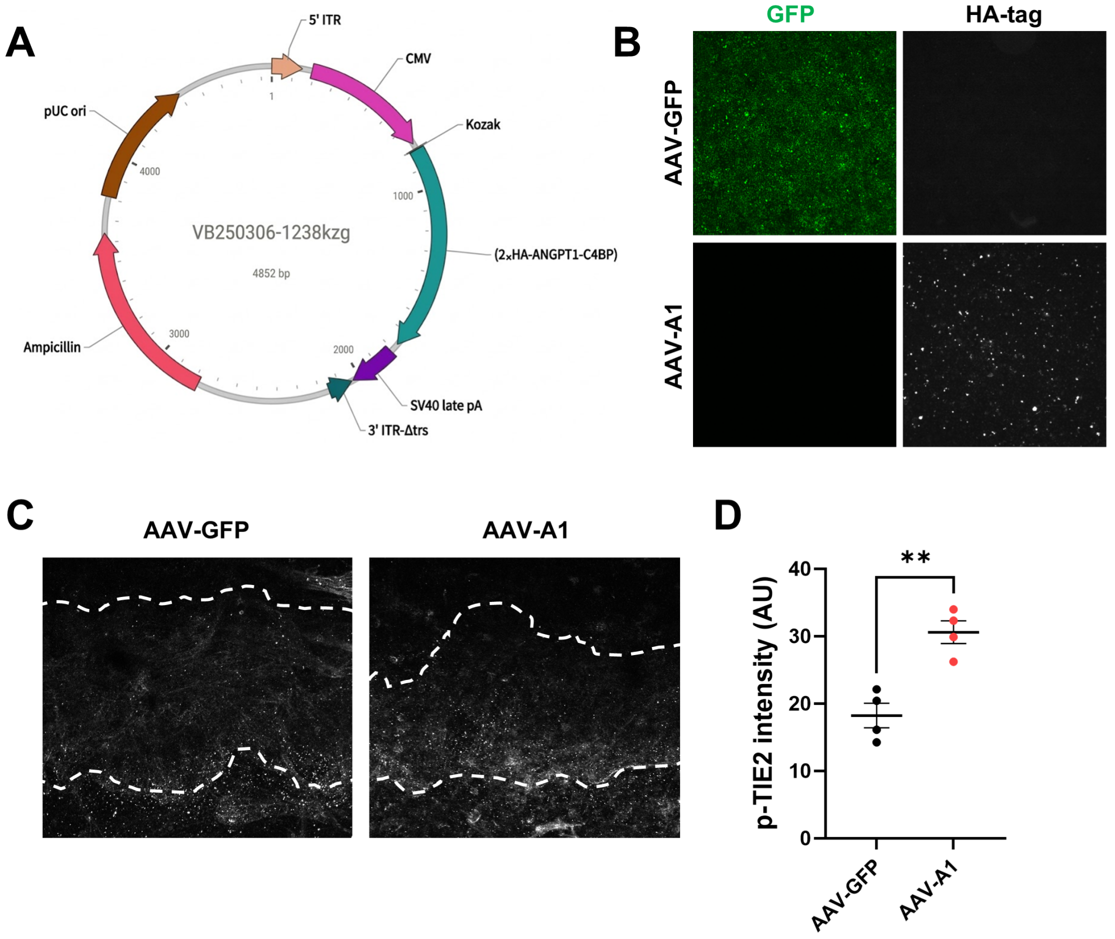


**Supplementary Figure S9. Design, in vitro validation, and in vivo p-TIE2 activation by AAV-heptaAng1.**

(A) Schematic of the self-complementary AAV expression construct encoding 2×HA-ANGPT1-C4BP under the control of the CMV promoter. (B) Representative fluorescence images of HEK293 cells exposed to control AAV-GFP or AAV-heptaAng1 (AAV-A1). GFP signal was detected only in the AAV-GFP condition, whereas HA-tag signal was detected only in the AAV-A1 condition, confirming in vitro expression of the AAV-encoded 2×HA-tagged

ANGPT1-C4BP transgene. (C) Representative p-TIE2 immunostaining images of the Schlemm's canal (SC) region from older mice treated with AAV-GFP or AAV-A1. Dashed lines outline the SC region. (D) Quantification of p-TIE2 intensity in the SC region, showing increased p-TIE2 in AAV-A1-treated eyes. Data are presented as mean ± SEM. Each dot represents one eye. Two-tailed unpaired Student's t-test was used. **$P < 0.01$.

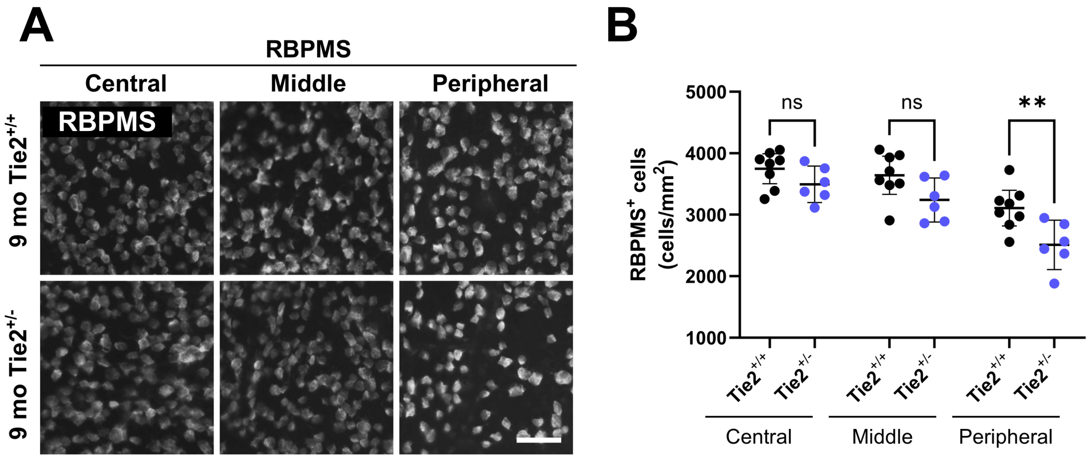


**Supplementary Figure S10.** ***Tie2*** **haploinsufficiency reduces peripheral retinal ganglion cell density at 9 months.**

(A) Representative RBPMS immunostaining images from central, middle, and peripheral retinal regions of 9-month-old *Tie2*$^{+/+}$ and *Tie2*$^{+/-}$ mice. Scale bar, 50 μm. (B) Quantification of RBPMS$^{+}$ retinal ganglion cell density in the indicated retinal regions. Data are presented as mean ± SEM. Two-tailed unpaired Student's t-tests were used for each retinal region. ns, not significant; $^{**}P < 0.01$.

**Supplementary Table S1.**

**Human donor age and death-to-preservation interval**

| Age | Death-to-preservation |
|---|---|
| 17 | 6 h 44 min |
| 29 | 1 h 01 min |
| 48 | 11 h 07 min |
| 65 | 13 h 44 min |
| 67 | 6 h 10 min |
| 68 | 11 h 23 min |
| 74 | 10 h 17 min |
| 72 | 13 h 59 min |

**Supplementary Table S2.**

**Human donor age and death-to-preservation interval**

| Age | Death-to-preservation |
|---|---|
| 48 | 10 h 25 min |
| 72 | 13 h 59 min |
| 58 | 13 h 12 min |
| 41 | 4 h 57 min |
| 24 | 7 h 38 min |
| 48 | 9 h 35 min |
| 40 | 13 h 55 min |
| 62 | 15 h 05 min |

### Supplementary Table S3.
### ssODN sequences used for loxP insertion at the *Vegfa* locus

| **Name** | **Sequence (5′–3′)** |
|---|---|
| *Vegfa*_loxP_ssODN1 | ggactggcaggtgtctctcgaagcttctgtgtctgtgcatgatatcagtgtcagaaggcatagccaacatagctc<br>cttctaactagttcttactagtattgtggcctctgATAACTTCGTATAATGTATGCTATACGAAGT<br>TATcccgggggatgaatggtggtgtttctctgagtctctagatctccat |
| *Vegfa*_loxP_ssODN2 | actagcgtccacagcagagtgcaggagagcgagtgggtgccacagATAACTTCGTATAATGTAT<br>GCTATACGAAGTTATcccgggggcagtttctgtgtatcaggggatggacgtagcctgggccttgttggg<br>tggtgactgtggccatccaggtttttaggtcccaggagatttcttggttgtgtgtgtatgcctttccgtagggctgtg |

Lowercase letters indicate homology arms, and uppercase letters indicate the loxP sequence.

**Supplementary Table S4.**

**Materials used in the present study**

| Category | Reagent / item | Company / source | Identifier |
|---|---|---|---|
| Primary antibody | Anti-PECAM-1 (CD31) | BD Biosciences | 553370 |
| Primary antibody | Anti-Iba1 | Wako | 019-19741 |
| Primary antibody | Anti-CD45 | R&D Systems | AF114-SP |
| Primary antibody | Anti-LYVE1 | AngioBio / Labome | 11-034 |
| Primary antibody | Anti-CD68 | Abcam | ab125212 |
| Primary antibody | Anti-RBPMS | PhosphoSolutions | 1832-RBPMS |
| Primary antibody | Anti-Phospho-TIE2 | R&D Systems | AF2720 |
| Primary antibody | Anti-HA | Cell Signaling Technology | 3724S |
| RNAscope probe | Probe - Mm-*Vegfa* | ACD; Bio-Techne | 312931 |
| RNAscope probe | Probe - Hs-*VEGFA* | ACD; Bio-Techne | 423161 |
